# J-domain proteins stimulate PKL-mediated chromatin compaction at H3K4-hypomethylated genomic loci

**DOI:** 10.64898/2026.09.14.751395

**Authors:** Bin-Bin Guan, Zijuan Yan, Ping Du, Youyang Sia, Dan-Yang Yuan, Yu-Xi Luo, Jia-Xin Li, Xiao-Min Su, Yin-Na Su, Jing Guo, Zhen-Zhen Liu, Xin-Yi Liu, Lin Li, She Chen, Zhucheng Chen, Xin-Jian He

## Abstract

Chromatin accessibility varies widely across distinct genomic regions in eukaryotes, yet the mechanisms governing these differential patterns remain poorly understood. Here, we identify a subfamily of functionally redundant J-domain proteins (JDPs) that assemble into a protein complex with PICKLE (PKL), an evolutionarily conserved CHD3-type chromatin remodeler, in *Arabidopsis thaliana*. JDPs are required not only for maintaining PKL protein stability but also for stimulating its nucleosome remodeling and ATPase activities. A previously uncharacterized histone-binding domain (HBD) within JDPs specifically recognizes the N-terminal tail of histone H3 when the H3K4me3 modification is absent. This interaction enhances PKL-mediated nucleosome sliding and ATPase activities *in vitro*, and promotes PKL-dependent chromatin compaction at H3K4me3-depleted genomic loci *in vivo*. The PKL-JDP complex drives chromatin compaction to repress developmentally regulated genes, thereby governing key developmental phase transitions, including the embryo-to-seedling transition and flowering. Collectively, these findings uncover a distinct mechanism by which the absence of H3K4me3 is sensed to initiate regional chromatin compaction, repress developmentally regulated genes, and facilitate key developmental transitions.

## Introduction

The packaging of genomic DNA into chromatin imposes a fundamental barrier to all DNA-templated processes, with transcription requiring precise and dynamic regulation of chromatin accessibility.^1^ Genome-wide profiling of chromatin accessibility has revealed a highly non-uniform landscape: accessibility is typically enriched at transcription start site (TSS)-flanking regions, where transcriptional initiation occurs, but is markedly reduced in intergenic regions and heterochromatin domains.^2–5^ Beyond this static landscape, chromatin accessibility is dynamically regulated, enabling precise spatiotemporal control of gene expression.^6,7^ Dynamic regulation of chromatin accessibility coordinates diverse developmental processes in both plants and metazoans.^8–11^ The establishment and maintenance of appropriate accessibility patterns are therefore fundamental to developmental programming, yet the underlying mechanisms remain incompletely understood.

ATP-dependent chromatin remodelers play a central role in shaping chromatin accessibility by utilizing ATP hydrolysis to reposition, evict, or modify nucleosomes.^12–14^ Among the four major families (SWI/SNF, ISWI, CHD, and INO80) of chromatin remodelers, the CHD family, particularly the CHD3 subfamily, is distinguished by its predominant function in chromatin compaction and transcriptional repression.^14^ In metazoans, CHD3-type remodelers (such as Mi-2/CHD3 and CHD4) serve as the core ATPase subunits of the NuRD (nucleosome remodeling and deacetylase) complex, which represses gene expression by promoting nucleosome compaction and facilitating histone deacetylation.^15–17^ These activities are essential for coordinating stem cell maintenance and differentiation, thereby governing a wide range of developmental processes.^18,19^ The targeting of CHD3 remodelers to specific genomic loci is achieved through interactions with associated factors, including methyl-CpG-binding proteins (MBD2/3), metastasis-associated proteins (MTA1/2/3), and sequence-specific transcription factors, enabling context-dependent repression.^20–22^

In *Arabidopsis thaliana*, the CHD3-type remodeler PICKLE (PKL) is a well-established regulator of multiple developmental processes, including the embryonic-to-seedling transition and flowering.^23–31^ During germination, PKL represses embryonic identity genes by promoting the deposition of the repressive histone mark H3K27me3, a function that requires its interaction with Polycomb Repressive Complex 2 (PRC2).^28,32,33^ PKL also regulates flowering time by modulating the expression of flowering regulatory genes.^29,34^ Additionally, PKL contributes to RNA-directed DNA methylation (RdDM), facilitating the silencing of transposable elements and transgenes.^35^ Notably, PKL targets both H3K27me3-marked regions and DNA-methylated regions, two classes of repressive chromatin that are largely mutually exclusive in the Arabidopsis genome.^36–38^ This observation raises a fundamental question: how does a single chromatin remodeler selectively associate with two distinct repressive chromatin environments?

Unlike their metazoan counterparts, which assemble into multi-subunit NuRD/Mi-2 complexes,^16,20^ PKL was long thought to function as a monomer.^39,40^ Although PKL interacts with the histone deacetylases HDA6 and HDA9 and the H3K27 methyltransferase CLF, these interactions are unlikely to form stable protein complexes.^41,42^ Moreover, the tandem plant homeodomain (PHD) fingers found in the N-terminal region of metazoan CHD3 family remodelers can facilitate its association with repressive chromatin,^43,44^ whereas PKL possesses only a single PHD finger, and this domain is dispensable for PKL function *in vivo*.^33,45^ The transcriptional repressors VAL1 and VAL2 can recruit PKL to specific genomic loci, but their loss only modestly affects PKL recruitment a subset of its target loci.^30^ These observations suggest that PKL employs a previously uncharacterized alternative mechanism to achieve regional specificity in its chromatin compaction activity.

Here, we identify four functionally redundant J-domain proteins (JDP1-4) that form a stable protein complex with PKL. We demonstrate that the PKL-JDP complex preferentially compacts chromatin at distal promoter and intergenic regions that are typically depleted of the active histone mark H3K4me3. JDPs contain a histone-binding domain (HBD) at their C-termini, which selectively binds the histone H3 tail in an H3K4 hypomethylation-dependent manner. This HBD is required for stimulating PKL-mediated nucleosome remodeling *in vitro* and for enforcing PKL-dependent restriction of chromatin accessibility at target genomic loci *in vivo*. Our findings support a model in which the PKL-JDP complex senses the absence of H3K4me3 to initiate chromatin compaction, thereby enabling the subsequent deposition of repressive marks such as H3K27me3 and DNA methylation.

## Results

### PKL interacts with J-domain proteins to form a protein complex

To identify proteins interacting with PKL, we generated transgenic Arabidopsis plants expressing a native promoter-driven PKL transgene fused to an N-terminal FLAG tag. Affinity purification coupled with mass spectrometry (AP-MS) using an anti-FLAG antibody revealed three PKL-interacting proteins: two J-domain proteins encoded by tandem genes with identical coding sequences (AT2G05250 and AT2G05230), which we designated J-domain protein 1 and 2 (JDP1 and JDP2), and a small uncharacterized protein (AT1G01730), which we named PKL-associated protein (PAP) (Fig. 1a and Supplementary Data 1).

**Fig. 1.**
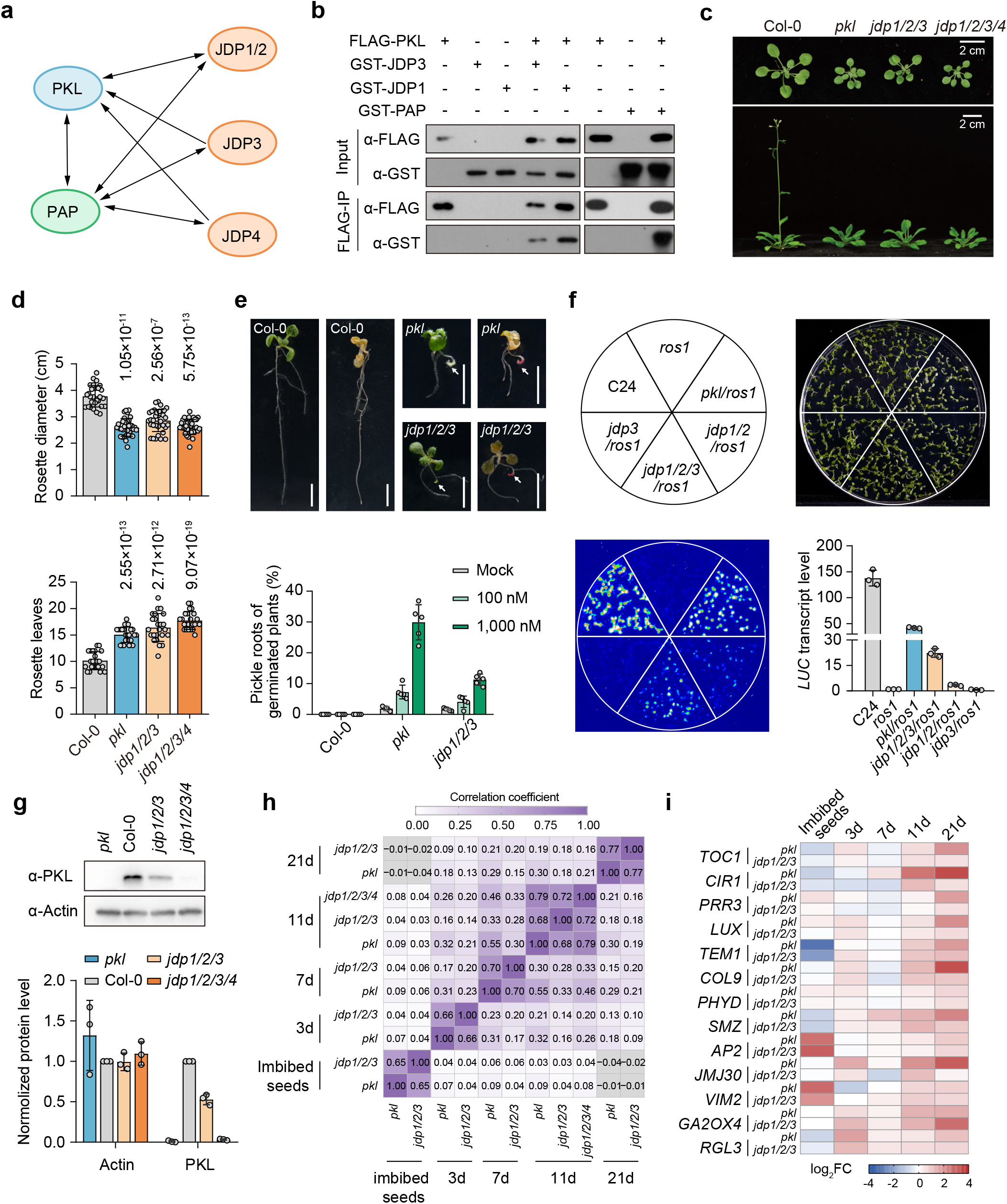
| J-domain proteins interact with PKL to co-regulate plant development and transcription. (**a**) Interactions of PKL, JDPs and PAP proteins determined by AP–MS. Arrow direction indicates the prey proteins identified for each bait. (**b**) Confirmation of direct interactions between PKL and JDP1, JDP3, or PAP via *in vitro* pull-down assays. Recombinant FLAG- or GST-tagged PKL, JDP1, JDP3, and PAP proteins were expressed *in vitro*, purified, and subjected to pull-down analyses. (**c, d**) Impacts of *pkl*, *jdp1/2/3*, and *jdp1/2/3/4* on plant development and flowering time. In (**c**), Vegetative developmental phenotypes are shown for 3-week-old plants (top), and flowering-time phenotypes are shown for 5-week-old plants (bottom). In (**d**), quantification of rosette width (left) of 3-week-old plants and rosette leaf number (right) at bolting is shown. Circles represent the values of indicated samples. Data are presented as mean ± standard deviation (SD) and *p* values were determined by two-tailed Student’s t-test. (**e**) Pickle root morphology and quantitative analysis in 10-day-old seedlings. Representative images of Col-0 (wild type), *pkl*, and *jdp1/2/3* mutant seedlings before (left) and after (right) Sudan Red staining are shown in the top panel. White arrows indicate pickle roots. Scale bar, 0.5 cm. Quantification of pickle root penetrance in seedlings grown on MS medium with or without uniconazole-P treatment are shown in the bottom panel. Data are presented as mean ± SD, with circles representing individual biological replicates. Over 80 geminated seedlings are quantified per replicate. (**f**) Effects of *pkl*, *jdp1/2*, *jdp3*, and *jdp1/2/3* mutations on the silencing of the *RD29A-LUC* transgene. Luminescence imaging and quantification of *LUC* transcription levels in indicated mutants are shown. All mutants are derived from the ecotype C24. Data are presented as mean ± SD, with individual data points represented by circles. (**g**) PKL protein abundance in Col-0, *pkl*, *jdp1/2/3* and *jdp1/2/3/4* plants. Immunoblot images (the top panel) and quantification (the bottom panel) of the PKL protein are shown. Actin was detected as an internal loading control. Data are presented as mean ±SD from three independent biological replicates. (**h**) Pairwise correlation coefficients of expression changes (log_2_ fold change) for differentially expressed genes in *pkl* and *jdp* mutants relative to wild-type plants during seedling development. The color bar shows the relationship between color shades and correlation values, with coefficients shown in each heatmap cell. (**i**) Heatmap showing the expression changes (log_2_ fold change) of representative flowering repressors in *pkl* and *jdp1/2/3* mutants compared to wild-type plants.

To validate these interactions, we generated transgenic plants expressing FLAG-tagged JDP1 and PAP and performed AP-MS with an anti-FLAG antibody. This analysis not only confirmed the interactions between PKL and JDP1/2 and between PKL and PAP, but also identified two additional JDP paralogs, termed J-domain protein 3 and 4 (JDP3 and JDP4), that interact with PAP (Fig. 1a and Supplementary Fig. 1a). We subsequently expressed FLAG-tagged JDP3 and JDP4 in Arabidopsis plants and further confirmed their interactions with PKL by AP-MS (Fig. 1a). These results demonstrate that PKL interacts with four JDP paralogs (hereafter collectively referred to as JDPs) and PAP. Notably, AP-MS did not detect interactions among the JDP paralogs themselves, suggesting that JDPs function as mutually-exclusive subunits within the complex. *In vitro* pull-down assays further showed that PKL directly binds JDP1, JDP3 and PAP (Fig. 1b), supporting the conclusion that PKL assembles into a multi-protein complex comprising JDPs and PAP. In addition, AP-MS data revealed that ATRX predominantly associates with the PKL-JDP3-PAP subcomplex, indicating that ATRX is not a stable, universal constituent of PKL-containing complexes (Supplementary Fig. 1a, b).

### J-domain proteins and PKL co-regulate development and flowering time

Previous studies have demonstrated that loss-of-function mutations in *PKL* cause multiple developmental defects in Arabidopsis.^23,34,35^ To investigate whether JDPs and PAP function together with PKL in regulating these phenotypic traits, we generated *jdp* single, double, triple, and quadruple mutants, as well as a *pap* single mutant, using the CRISPR-Cas9-mediated gene editing system. All mutant alleles harbored 1–2 bp insertions or deletions that caused frameshift mutations in the corresponding target genes (Extended Data Fig. 1a). The *jdp1 jdp2* (*jdp1/2*) double mutant displayed reduced plant size and delayed flowering, whereas the *jdp3* single mutant did not exhibit these phenotypes (Extended Data Fig. 1b, c).

The developmental defects observed in the *jdp1/2* double mutant were further exacerbated in the *jdp1 jdp2 jdp3* (*jdp1/2/3*) triple mutant and became even more pronounced in the *jdp1 jdp2 jdp3 jdp4* (*jdp1/2/3/4*) quadruple mutant (Fig. 1c, d, and Extended Data Fig. 1a-c). Notably, combining *jdp* mutations with the *pkl* mutation did not result in additive phenotypic effects (Extended Data Fig. 1b, c), indicating that JDPs and PKL function within a shared molecular pathway. In contrast, the *pap* mutant exhibited no detectable changes in plant size or flowering time in the wild-type, *pkl*, or *jdp* mutant backgrounds (Extended Data Fig. 1b, c). These results suggest that JDPs cooperate with PKL to regulate plant development and flowering time, whereas PAP is dispensable for these developmental processes in Arabidopsis. We therefore focused subsequent analyses on JDPs within the PKL-JDP complex.

Because the *pkl* mutant roots exhibit embryonic “pickle” traits with partial penetrance,^23,46^ we next examined whether JDPs are involved in this developmental trait. While wild-type plants showed no embryonic root phenotypes, approximately 2% of both *pkl* and *jdp1/2/3* triple mutant plants displayed embryonic root traits. The penetrance of this phenotype increased in a dose-dependent manner upon treatment with uniconazole-P, a specific inhibitor of gibberellin (GA) biosynthesis (Fig. 1e). These findings indicate that JDPs and PKL act synergistically to regulate the developmental transition from embryogenesis to postembryonic root morphogenesis.

PKL has been shown to mediate transcriptional silencing of an *RD29A* promoter-driven luciferase reporter (*RD29A-LUC*) transgene in the *ros1* DNA demethylation mutant background.^35^ To investigate whether JDPs also participate in this silencing process, we introduced *pkl* and *jdp* mutations into the *ros1* mutant carrying the *RD29A-LUC* transgene (Fig. 1f). Consistent with the previous reports,^35^ the *pkl* mutation relieved silencing of the *RD29A-LUC* transgene (Fig. 1f). Transgene silencing was derepressed in the *jdp1/2* double mutant but not in the *jdp3* single mutant, and this effect was further enhanced in the *jdp1/2/3* triple mutant (Fig. 1f). These results demonstrate that JDPs cooperate with PKL to mediate transgene silencing in Arabidopsis.

Notably, PKL protein levels were markedly reduced in both the *jdp1/2/3* triple and *jdp1/2/3/4* quadruple mutants (Fig. 1g). This reduction was more severe in the quadruple mutant, which correlated with its more pronounced developmental defects. These findings suggest that JDPs regulate plant development, at least in part, by maintaining PKL protein abundance.

### PKL and JDPs co-regulate genome-wide transcription

To determine whether JDPs and PKL co-regulate genome-wide transcription, we performed RNA sequencing (RNA-seq) on *jdp1/2/3*, *pkl*, and wild-type plants at multiple developmental stages: imbibed seeds, and 3-, 7-, 11-, and 21-day-old plants. RNA-seq analysis revealed a strong positive correlation in protein-coding gene (PCG) expression changes among these mutants at the same developmental stages (Fig. 1h and Supplementary Data 2). Gene Ontology (GO) analysis indicated that differentially expressed genes (DEGs) in *jdp1/2/3* and *pkl* mutants were enriched in a wide range of biological processes. We focused on GO terms associated with embryo development and flowering time and analyzed DEG enrichment within these categories (Extended Data Fig. 2a). Notably, many genes involved in seed oil body biogenesis and lipid storage were differentially expressed in both mutants. Given that *pkl* mutants have been reported to accumulate seed-like fatty acids and oil bodies in the root system,^23^ this suggests that the PKL-interacting JDPs participate in these processes together with PKL.

Furthermore, the expression levels of numerous flowering repressors were increased in both *pkl* and *jdp1/2/3* mutants, particularly in 11- and 21-day-old plants (Fig. 1i). These included circadian rhythm genes (*TOC1*, *CIR1*, *PRR3*, and *LUX*), photoperiodism-related genes (*TEM1*, *COL9*, *PHYD* and *SMZ*), and other flowering repressors.^47–54^ Consistently, several flowering-promoting genes were correspondingly downregulated (Extended Data Fig. 2b). These expression changes likely contribute to the late-flowering phenotype observed in these mutants. In addition, *GA2OX4* and *RGL3*, which act as repressors of GA signaling, were upregulated in 3-day-old mutants.^55,56^ As GA signaling promotes the embryo-to-seedling transition, their increased expression may contribute to the enhanced penetrance of embryonic traits in the roots of *pkl* and *jdp1/2/3* mutants.

We also performed RNA-seq on 11-day-old *jdp1/2/3/4* quadruple mutants, and found that the number of DEGs in *jdp1/2/3/4* mutants was increased compared to *jdp1/2/3* mutants and was comparable to that in *pkl* mutants, with substantial overlap among the three DEG sets (Extended Data Fig. 2c, d). In all three mutants, upregulated DEGs consistently outnumbered downregulated DEGs (Extended Data Fig. 2d). Moreover, the gene expression profile of *pkl* mutants more closely resembled that of *jdp1/2/3/4* mutants than that of *jdp1/2/3* mutants (Extended Data Fig. 2e), further supporting the conclusion that the four JDP paralogs function redundantly with PKL to co-regulate gene expression.

Given that PKL has been shown to mediate transcriptional silencing of transposable elements (TEs),^35^ we next examined TE expression in the *pkl* and *jdp1/2/3/4* mutants. RNA-seq analysis identified numerous differentially expressed TEs in these mutants relative to wild type, with expression changes showing strong positive correlations (Extended Data Fig. 2f and Supplementary Data 3). Moreover, consistent with the pattern observed for PCGs, upregulated TEs were more prevalent than downregulated TEs in all three mutants (Supplementary Data 3). These results support a model in which JDPs and PKL function cooperatively within the PKL-JDP complex to mediate transcriptional repression of both PCGs and TEs.

### PKL and JDPs co-target genomic loci to restrict chromatin accessibility

To determine whether PKL and JDPs share common genomic targets, we performed chromatin immunoprecipitation followed by sequencing (ChIP-seq) using transgenic Arabidopsis plants expressing GFP-tagged PKL and JDP1. ChIP-seq analysis identified 20,247 PKL binding peaks and 19,348 JDP1 binding peaks (Fig. 2a). Assigning these peaks to their nearest genes revealed 20,043 PKL target genes and 19,015 JDP1 target genes, with 18,383 genes co-targeted by both factors (Fig. 2b and Supplementary Data 4). PKL and JDP1 binding signals exhibited a strong positive correlation across PKL target genes and were prominently enriched in regions flanking the transcription start site (TSS) (Fig. 2c). These results indicate that PKL and JDP1 extensively co-occupy TSS-proximal regions throughout the genome.

**Fig. 2.**
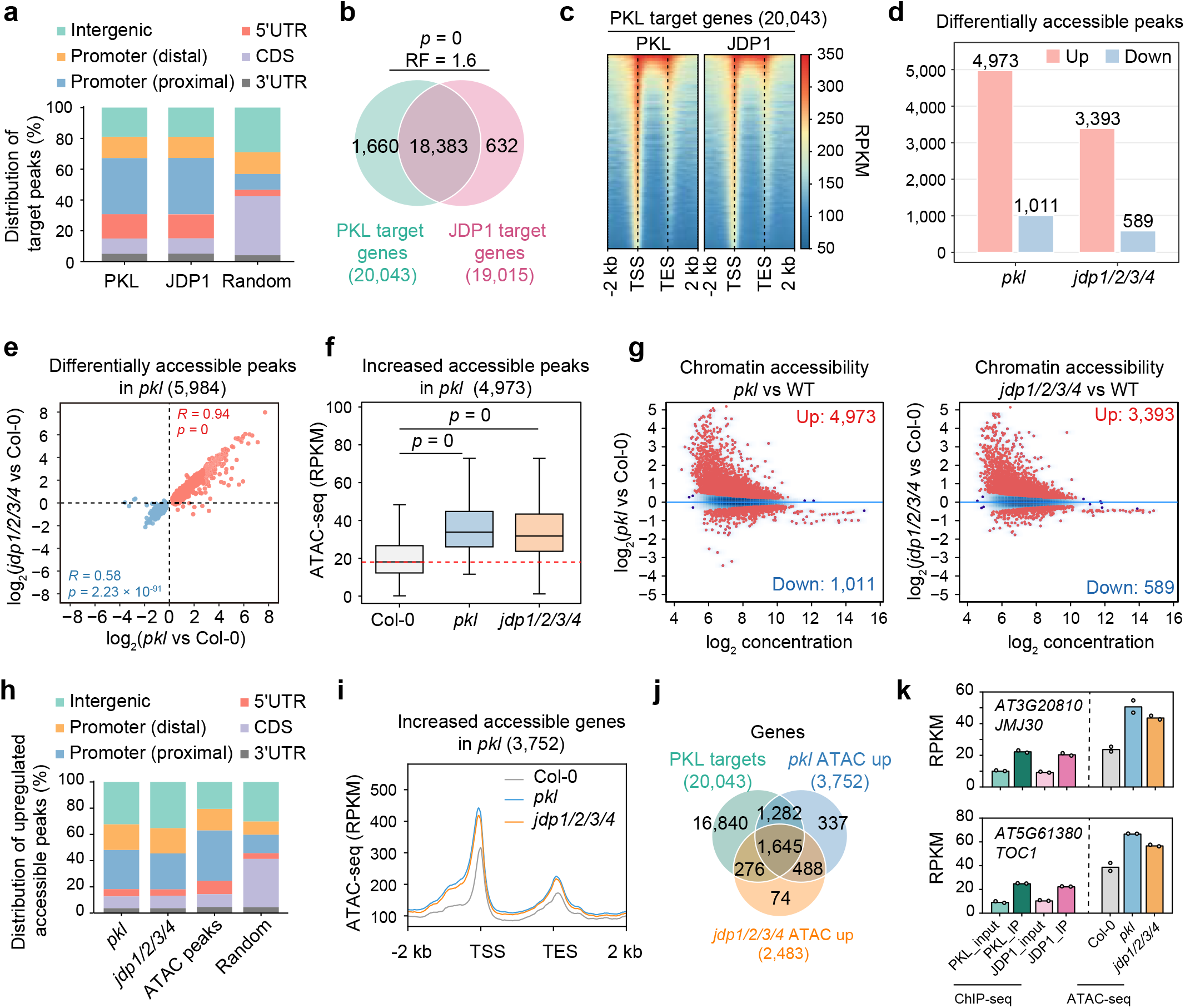
| Co-repression of chromatin accessibility by PKL and JDPs at low accessible regions. (**a**) Genomic annotation of PKL and JDP1 ChIP-seq peaks. Distal promotor: 401–1,000 bp upstream of transcription start site (TSS); proximal promotor: 1–400 bp upstream of TSS. (**b**) Overlap between PKL target genes and JDP1 target genes. The significance of the overlap is indicated by *p* values and representation factor (RF); *p* values were determined using a one-tailed hypergeometric test. (**c**) Heatmap showing the ChIP-seq signals of PKL and JDP1 at PKL target genes. Target genes are sorted by PKL enrichment intensity. TSS, transcription start site; TES, transcription end site. (**d**) Number of genomic loci with upregulated and downregulated chromatin accessibility in *pkl* and *jdp1/2/3/4* mutants relative to wild-type plants (FDR < 0.05). (**e**) Correlation of chromatin accessibility changes in *pkl* and *jdp1/2/3/4* mutants compared to wild-type Col-0 plants. Analyses were performed on genomic loci with significantly altered accessibility in the *pkl* mutant (FDR < 0.05). (**f**) Boxplots showing the effects of *pkl* and *jdp1/2/3/4* mutations on chromatin accessibility. Analyses were performed on genomic loci with significantly upregulated accessibility in the *pkl* mutant (FDR < 0.05). *p* values were determined using a two-tailed paired Mann-Whitney U test for non-normally distributed data. Center lines and box edges indicate medians and interquartile ranges (IQR), respectively; whiskers extend to within 1.5×the IQR. (**g**) Scatter plots showing genomic loci with altered chromatin accessibility in *pkl* and *jdp1/2/3/4* mutants compared to wild-type plants. Red dots represent genomic loci with significant changes in chromatin accessibility (FDR < 0.05). (**h**) Genomic annotation of loci with significantly upregulated accessibility (FDR <0.05) in *pkl* and *jdp1/2/3/4* mutants. Distal promotor: 401–1,000 bp upstream of transcription start site (TSS); proximal promotor: 1–400 bp upstream of TSS. The distribution of ATAC peaks represents the total chromatin accessible peaks in Col-0 and *pkl*. (**i**) Metaplots showing ATAC-seq signal patterns in wild type, *pkl*, and *jdp1/2/3/4* mutant plants. Analyses were performed using genes with significantly upregulated accessibility in the *pkl* mutant (FDR < 0.05). (**j**) Overlap between PKL target genes and genes with up-regulated ATAC-seq signals in *pkl*, and *jdp1/2/3/4* mutants. The significance of the overlap is indicated by *p* values and representation factors (RF). PKL targets vs *pkl* ATAC up: *p* = 5.05×10-115, RF = 1.3; PKL targets vs *jdp1/2/3/4* ATAC up: *p* = 4.29×10-68, RF = 1.3; *pkl* ATAC up vs *jdp1/2/3/4* ATAC up: *p* = 0, RF = 7.5. *p* values were determined using a one-tailed hypergeometric test. (**k**) Quantitative analysis of PKL and JDP1 ChIP-seq peaks and ATAC-seq peaks with increased chromatin accessibility in *pkl* and *jdp1/2/3/4* mutants within the 2 kb region upstream of the TSS for representative flowering repressor genes.

Because PKL represses chromatin accessibility,^33^ we next performed Assay for Transposase-Accessible Chromatin using sequencing (ATAC-seq) in wild type, *pkl*, and *jdp1/2/3/4* mutants. Compared wild type, the *pkl* mutant exhibited 4,973 upregulated and 1,011 downregulated accessible chromatin peaks, whereas the *jdp1/2/3/4* mutant exhibited 3,393 upregulated and 589 downregulated peaks (Fig. 2d and Supplementary Data 5). In both mutants, increases in chromatin accessibility substantially outnumbered decreases. Moreover, changes in chromatin accessibility showed a strong positive correlation between the *pkl* and *jdp1/2/3/4* mutants (Fig. 2e), and the elevated accessibility observed in *pkl* was recapitulated in *jdp1/2/3/4* (Fig. 2f). These data support a cooperative role for PKL and JDPs in repressing chromatin accessibility.

Notably, the most strongly upregulated accessible peaks in both mutants were predominantly located at genomic loci with low basal accessibility in wild-type plants (Fig. 2g). Compared to the total set of ATAC-seq peaks, these upregulated accessible peaks were enriched in distal promoter and intergenic regions, which inherently exhibit lower accessibility than TSS-proximal regions (Fig. 2h). Metaplots further revealed that accessibility increases in the mutants extended from TSS-flanking regions into upstream distal promoter and intergenic regions, but not into downstream intragenic regions (Fig. 2i).

To assess whether the PKL-JDP complex directly restricts chromatin accessibility, we assigned upregulated ATAC-seq peaks to their nearest genes and intersected these genes with PKL target genes. We identified 3,752 and 2,483 genes with increased chromatin accessibility in the *pkl* and *jdp1/2/3/4*, respectively, and found that more than three-quarters of these genes were PKL targets (Fig. 2j). Among genes exhibiting increased chromatin accessibility in the *pkl* mutant, upregulated DEGs greatly outnumbered downregulated ones (Extended Data Fig. 3a), suggesting that the PKL-JDP complex mediates transcriptional repression by restricting chromatin accessibility. Consistent with this model, several key genes associated with flowering and embryo-to-seedling transition, which showed increased expression in *pkl* and *jdp* mutants, are direct targets of the PKL-JDP complex, with their chromatin accessibility increased in distal promoter regions in these mutants (Fig. 2k and Extended Data Fig. 3b). These results support a model in which the PKL-JDP complex directly restricts the spread of chromatin accessibility from TSS-flanking regions into distal upstream regions, thereby repressing the expression of target genes.

### Mapping the PKL-JDP1 interaction interface

To investigate the molecular basis of the PKL-JDP interaction, we generated a series of truncated JDP1 variants and assessed their ability to bind PKL using yeast two-hybrid (Y2H) and GST pull-down assays (Fig. 3a). Y2H analyses revealed that the JDP1 fragment spanning amino acids 301– 480 is both necessary and sufficient for PKL binding; we therefore designated this region as the PKL-interacting domain (PID) (Fig. 3b). Consistent with this result, GST pull-down assays confirmed that the PID mediates the interaction with PKL (Extended Data Fig. 4a).

**Fig. 3.**
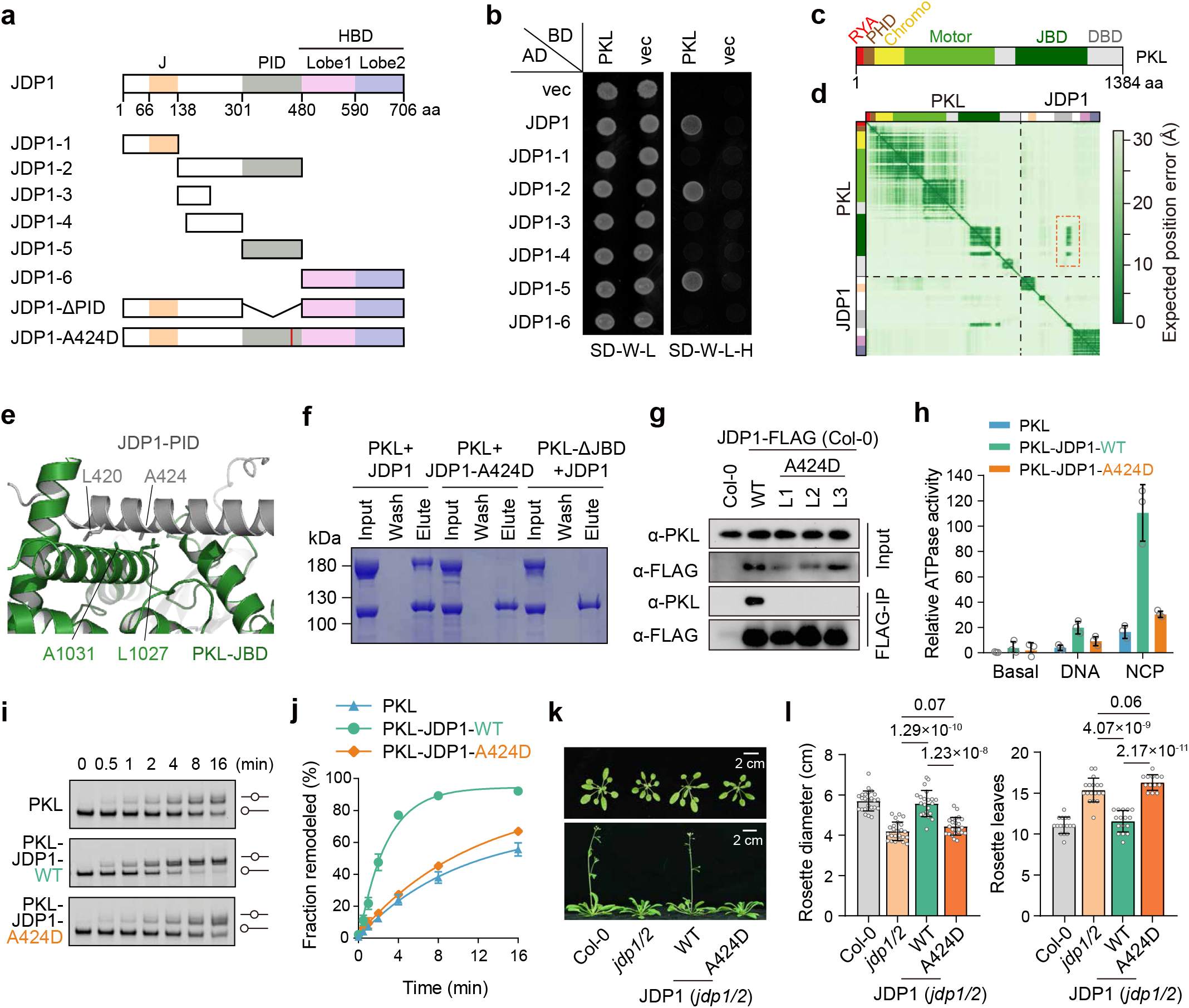
| JDP1 directly interacts with PKL and enhances its nucleosome-remodeling activity. (**a**) Schematic of JDP1 domain architecture. The PID (PKL-interacting domain) and HBD (histone binding domain) were identified in this study. A series of JDP1 truncated variants used in yeast two-hybrid and pull-down assays are shown. The A424D mutation site is marked by a red vertical line. (**b**) Yeast two-hybrid analysis of the PKL-JDP1 interaction. The interaction between PKL and full-length or truncated variants of JDP1 was assessed. Interaction was evaluated by yeast growth on synthetic dropout (SD) medium lacking tryptophan, leucine, and histidine (SD-W-L-H). (**c**) Schematic of PKL domain architecture. The RYA and JBD domains were identified in this study. (**d**) PAE matrix of the PKL-JDP1 complex generated by AlphaFold 3. (**e**) Predicted JDP1-PKL binding interface by AlphaFold3. The interacting residues are highlighted in stick representation. (**f**) GST pull-down assays showing the direct interaction between JDP1 and PKL. Truncation of the α-helical motif (residues 1023–1045) within the JBD and the JDP1-A424D mutation abolished the interaction. (**g**) Co-IP assay demonstrating that the A424D mutation disrupts the PKL-JDP1 interaction in Arabidopsis plants. (**h**) ATPase activities of PKL, PKL-JDP1-WT and PKL-JDP1-A424D. Data are presented as mean ±SD (n = 3). (**i, j**) Chromatin-remodeling activities of PKL, PKL-JDP1-WT and PKL-JDP1-A424D toward the 0N40 nucleosome. Representative gels are presented in (**i**), with quantification of the fractions of nucleosome slide shown in (**j**). Data are presented as mean ±SD (n = 3). The initial reaction rates (min^−1^) estimated by fitting and data are 0.07, 0.35 and 0.07 for PKL, PKL-JDP1-WT, and PKL-JDP1-A424D, respectively. (**k, l**) Effects of the JDP1-A424D mutation on plant development and flowering time. Representative morphological phenotypes of 3-week-old plants (top) and 5-week-old plants (bottom) are shown in (k); quantification of rosette width (left) in 3-week-old plants and rosette leaf number (right) in 5-week-old plants is shown in (l). Data are presented as mean ±SD. *p* values were determined using a two-tailed Student’s t-test. Individual data points are represented by circles.

PKL is a multi-domain chromatin remodeler, harboring a central ATPase motor domain, flanked by a conserved PHD finger and tandem chromo domains at the N-terminus, and a DNA-binding domain (DBD) at the C-terminus (Fig. 3c). To identify the JDP-binding region within PKL, we performed Y2H and GST pull-down assays using a panel of truncated PKL constructs (Extended Data Fig. 4b). These analyses demonstrated that the C-terminal half of PKL is required for interaction with JDPs (Extended Data Fig. 4c, d).

AlphaFold3 predicted that a previously uncharacterized C-terminal region of PKL (residues 825-1200) directly interacts with the PID of JDP1, with inter-chain PAE < 10 Å and interface residue pLDDT (predicted Local Distance Difference Test) > 70 (Fig. 3c-e). We designated this region as the JDP1-binding domain (JBD). A high pairwise ipTM (interface predicted Template Modeling score) of 0.66 was generated for this interaction prediction. The JBD of PKL contains a domain of unknown function 1087 (DUF1087) and a SANT-like domain, both of which are conserved not only in plants but also in metazoans (Supplementary Fig. 2). Within the JDB-PID interaction interface, JDP1-A424 packs against the hydrophobic side chains of PKL A1031 and L1027 (Fig. 3e). Consistent with this prediction, A424 is conserved among JDP1 homologs across higher plant lineages, including angiosperms and gymnosperms (Extended Data Fig. 5a).

To experimentally validate the functional importance of the JBD-PID interaction, we disrupted the predicted interface using two complementary approaches: (1) deletion of a conserved α-helix (residues 1023-1045) within the JBD of PKL, and (2) introduction of a site-specific A424D substitution in the PID of JDP1. Both perturbations completely abolished the PKL-JDP1 interaction, as demonstrated by *in vitro* pull-down assays (Fig. 3f). This result was further corroborated by co-IP assays of endogenous proteins in Arabidopsis (Fig. 3g). These findings demonstrate that PKL engages JDP1 through a specific JBD-PID interaction both *in vitro* and *in vivo*.

### JDP1 enhances the chromatin-remodeling activity of PKL

Previous *in vitro* analyses revealed that PKL alone possesses chromatin-remodeling activity.^39^ We next tested the impact of the assembly of the PKL-JDP complex on PKL’s enzymatic functions. Consistent with prior reports,^39^ PKL’s ATPase activity was modestly stimulated by DNA and strongly activated by nucleosomes (Fig. 3h). Addition of JDP1 further enhanced the ATPase activity, with a more pronounced effect observed in the presence of nucleosomes (Fig. 3h), suggesting that JDP1 recognizes some histone-associated features within nucleosomes to potentiate PKL’s ATPase function.

We next assessed the effect of JDP1 on the nucleosome-remodeling activity of PKL using a nucleosome-centering assay. In this assay, end-positioned nucleosomes (0N40) are repositioned toward the center of the DNA fragment, resulting in slower-migrating bands on native gels. Consistent with the ATPase measurements, JDP1 enhanced the nucleosome-centering activity of PKL (Fig. 3i, j). In agreement with the dispensability of PAP for PKL-regulated developmental phenotypes (Extended Data Fig. 1b, c), PAP did not promote either the ATPase or nucleosome-remodeling activities of PKL (Extended Data Fig. 4e-g).

Given that the JDP1-A424D mutation disrupts the PKL-JDP1 interaction, we next assessed the activity of PKL in the presence of this mutant. In both ATPase and nucleosome-centering assays, JDP1-A424D failed to enhance PKL enzymatic or remodeling activities *in vitro* (Fig. 3h-j). Furthermore, we generated transgenic plants expressing JDP1-A424D in the *jdp1/2* mutant background. Unlike wild-type JDP1, the A424D variant did not rescue the developmental or flowering-time defects of *jdp1/2* mutants (Fig. 3k, l and Extended Data Fig. 5b). These results demonstrate that direct interaction with PKL is essential for JDP1 function both *in vitro* and *in vivo*.

JDP proteins contain a conserved N-terminal J domain (Fig. 3a and Extended Data Fig. 5c), which typically mediates interactions with HSP70 chaperones.^57,58^ Because the HPD tripeptide motif within the J domain is required for HSP70 binding, we asked whether this motif is necessary for JDP1 function in Arabidopsis. We therefore generated transgenic plants expressing JDP1-HPD/QQQ and JDP1-H/Q mutant variants in the *jdp1/2* background (Extended Data Fig. 5c). Unexpectedly, both variants fully rescued the developmental defects of *jdp1/2* mutants (Extended Data Fig. 5d, e). These results indicate that while JDPs are critical for PKL-dependent chromatin remodeling, their function in regulating development and flowering is independent of canonical HSP70 interactions.

### Structure of PKL bound to the nucleosome

To illustrate how JDPs contribute to the chromatin-remodeling function of PKL, we determined the structure of the PKL-JDP1 complex bound to the nucleosome by cryo-electron microscopy (cryo-EM) (Extended Data Fig. 6, 7 and Supplementary Table 1). The structure was determined at an overall resolution of 3.1 Å (Fig. 4a, b and Extended Data Fig. 7), which reveals the PKL-nucleosome interaction. The structure of JDP1 was undetected, probably due to the tethering of JDP1 to the PKL motor through a flexible spacer sequence (Fig. 3f). This structure reveals that the PKL-JDP complex engages the nucleosome mainly through the PKL motor. The structural observation was confirmed by cross-linking mass spectrometry (CL-MS), which provide support for the JBD-PID interaction between PKL and JDP1 (Extended Data Fig. 6). The CL-MS results also identified contacts with the histone proteins (more discussion below).

**Fig. 4.**
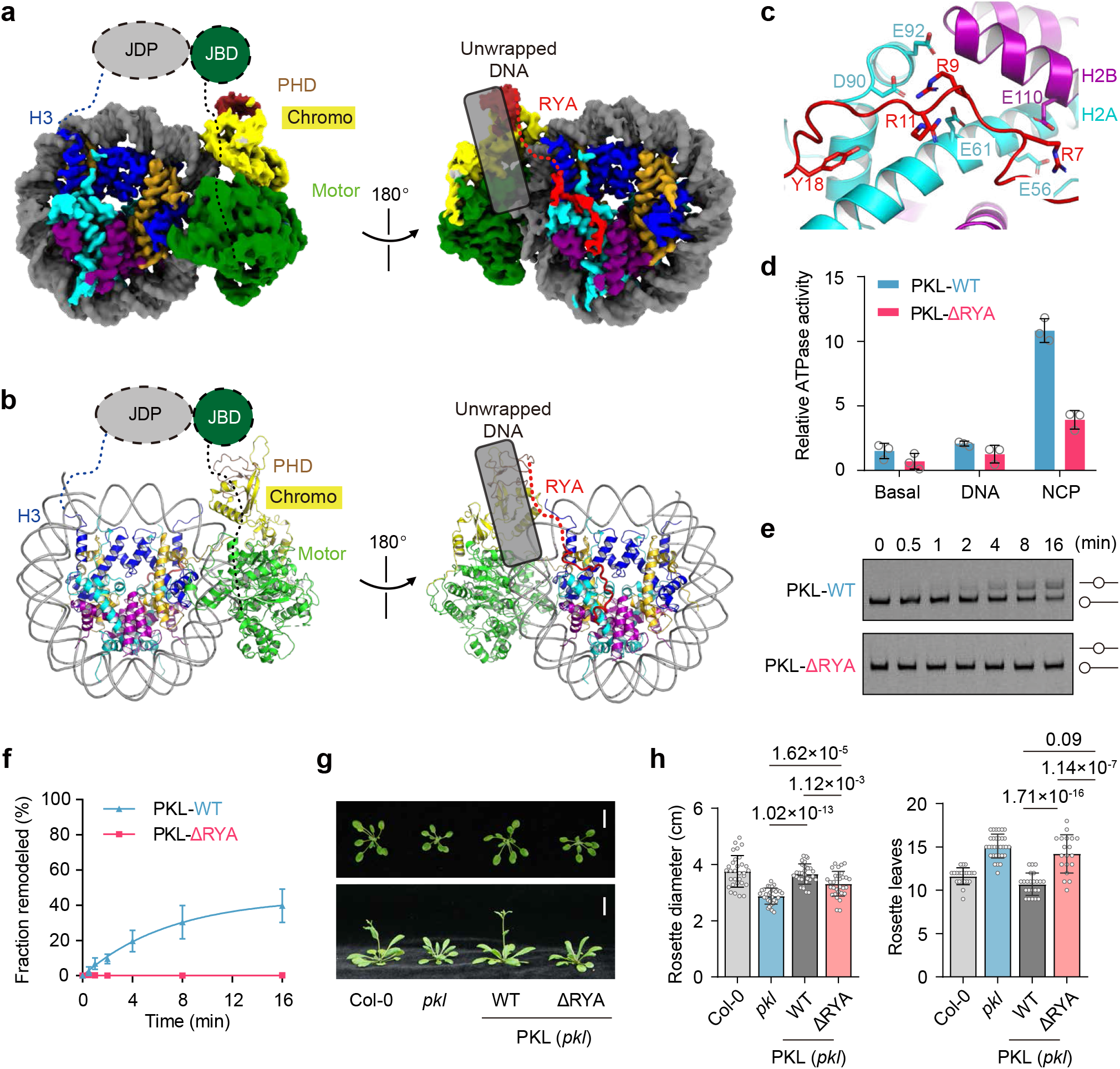
| A distinct role of the RYA motif revealed by the structure of PKL bound to the nucleosome. (**a**) Two views of the high-resolution composite maps of PKL bound to the nucleosome. The trajectory of unwrapped exit DNA is schematically illustrated as a box. (**b**) Two views of the ribbon model of PKL bound to the nucleosome. (**c**) Detailed interaction between the RYA motif of PKL and the acidic patch of the nucleosome. (**D**) Relative ATPase activities of PKL-WT and PKL-ΔRYA. Data are presented as mean ± SD (n = 3). (**e, f**) Chromatin-remodeling activities of PKL-WT and PKL-ΔRYA toward the 0N40 nucleosome. Representative gels are presented in (**e**), with quantification of the fractions of nucleosome slide shown in (**f**). Data are presented as mean ±SD (n = 3). The initial reaction rates (min^−1^) estimated by fitting and data are 0.07 and 0.00 for PKL-WT and PKL-ΔRYA, respectively. (**g, h**) Effects of RYA motif deletion on plant development and flowering time. Representative morphological phenotypes of 3-week-old plants (top) and 5-week-old plants (bottom) are shown in (**g**); quantification of rosette width (left) in 3-week-old plants and rosette leaf number (right) in 5-week-old plants is shown in (**h**). Data are presented as mean ±SD. *p* values were determined using a two-tailed Student’s t-test. Individual data points are represented by circles.

PKL engages the nucleosome in a manner largely similar to the known CHD proteins, including yeast Chd1 and human CHD4,^59,44^ Yet, different from its homologs, PKL induces a more subtle unwrapping conformation of the exit DNA (Fig. 4a, b). Chd1 drives dramatic exit DNA unwrapping, a process dependent on the DNA-binding ability of its DBD;^60^ in contrast, CHD4 did not induce DNA unwrapping, which is consistent with the absence of a stably associated DBD in its nucleosome-bound structure (Extended Data Fig. 7). PKL induces ∼10 bp of exit DNA unwrapping, despite lacking a stably bound DBD (Fig. 4a, b).

Interestingly, an arginine-tyrosine anchor (RYA, residues 4-25) motif at the N-terminus of PKL was identified to bind the nucleosome (Fig. 4C and Extended Data Fig. 7c). Similar to other chromatin remodelers,^61^ the motor domain of PKL binds to the nucleosome at super helical location 2 (SHL2). The RYA motif extends out of the N-terminal end of PHD, and reaches the H2A-H2B surface of the nucleosome disc opposite to the motor binding site. Arg7, Arg9, Arg11, and Tyr18 of the RYA motif recognize the acidic and adjacent hydrophobic surface of the nucleosome (Fig. 4c). The linker sequence between the RYA motif and PHD domain is expected to sterically clash with the exit DNA in the wrapped conformation, explaining the partially unwrapping conformation of the exit DNA (Fig. 4a). Notably, the RYA motif is conserved in Chd1 but absent from CHD4 (Extended Data Fig. 8). Therefore, PKL shows unique features distinct from the yeast and human homologs, with RYA recognizing the nucleosome, but without DBD to unwrap the exit DNA.

To investigate the functional significance of the RYA motif, we expressed a truncated PKL variant lacking this motif (PKL-ΔRYA), and evaluated its ATPase and remodeling activities. Deletion of the RYA motif abolished both ATPase and remodeling activities of PKL *in vitro* (Fig. 4d-f). Consistently, while transgenic expression of wild-type PKL fully rescued the reduced plant size and delayed flowering-time phenotypes of *pkl* mutants, PKL-ΔRYA failed to restore flowering time and only partially rescued plant size (Fig. 4g, h). These results demonstrate that PKL depends on its RYA motif to remodel nucleosomes and regulate plant development.

### The histone-binding domain of JDP1 preferentially recognizes hypomethylated H3K4

The mechanism of PKL activation by JDP1 is intriguing, as JDP1 achieves this without forming stable interactions with either the PKL motor domain or the nucleosome as shown by the cryo-EM structure. Remarkably, AlphaFold3 predicts that the HBD of JDP1 folds into two closely packed regions (Lobe1 and Lobe2) with unknown function, which nevertheless exhibit a topology reminiscent of the Bromo adjacent homology (BAH) domain (Supplementary Fig. 3a). Given that the canonical BAH domains are histone tail interactors,^62–67^ we investigated whether JDP1 directly engages histones. Pull-down assays demonstrated that JDP1 selectively interacts with histone H3, but not with H2A, H2B, or H4 (Supplementary Fig. 3b). Consistently, the CL-MS analysis further supported the interaction between JDP1 and H3 within the nucleosome-bound PKL-JDP1 complex (Extended Data Fig. 6c).

Importantly, the HBD of JDP1 specifically binds the N-terminal tail of H3 (residues 1-19), and this interaction is sensitive to the methylation status of H3K4 (Fig. 5a). H3K4me3 and, to a lesser extent, H3K4me2 markedly reduced the binding, whereas H3K9 methylation had no detectable effect on this interaction (Fig. 5a). Moreover, the HBD barely binds the internal segment of the H3 N-terminus (residues 21-44), regardless of the methylation status of H3K27 or H3K36. We next quantified the interaction using isothermal titration calorimetry (ITC). The analysis showed that the HBD exhibits a strong preference for hypomethylated H3K4, with comparable affinities for H3K4me0 and H3K4me1 (Fig. 5b). By contrast, no detectable binding to H3K4me2 and H3K4me3 was observed by ITC. The HBD is conserved across all four Arabidopsis JDP homologs and retained in JDP orthologs throughout plant evolution, from bryophytes to angiosperms (Supplementary Fig. 3c, d), indicating that HBD-containing J-domain proteins represents an evolutionarily conserved family in plants. Thus, our findings indicate that the previously unrecognized HBD in a subset of J-domain proteins mediate selective recognition of histone H3 tail in an H3K4 hypomethylation-dependent manner.

**Fig. 5.**
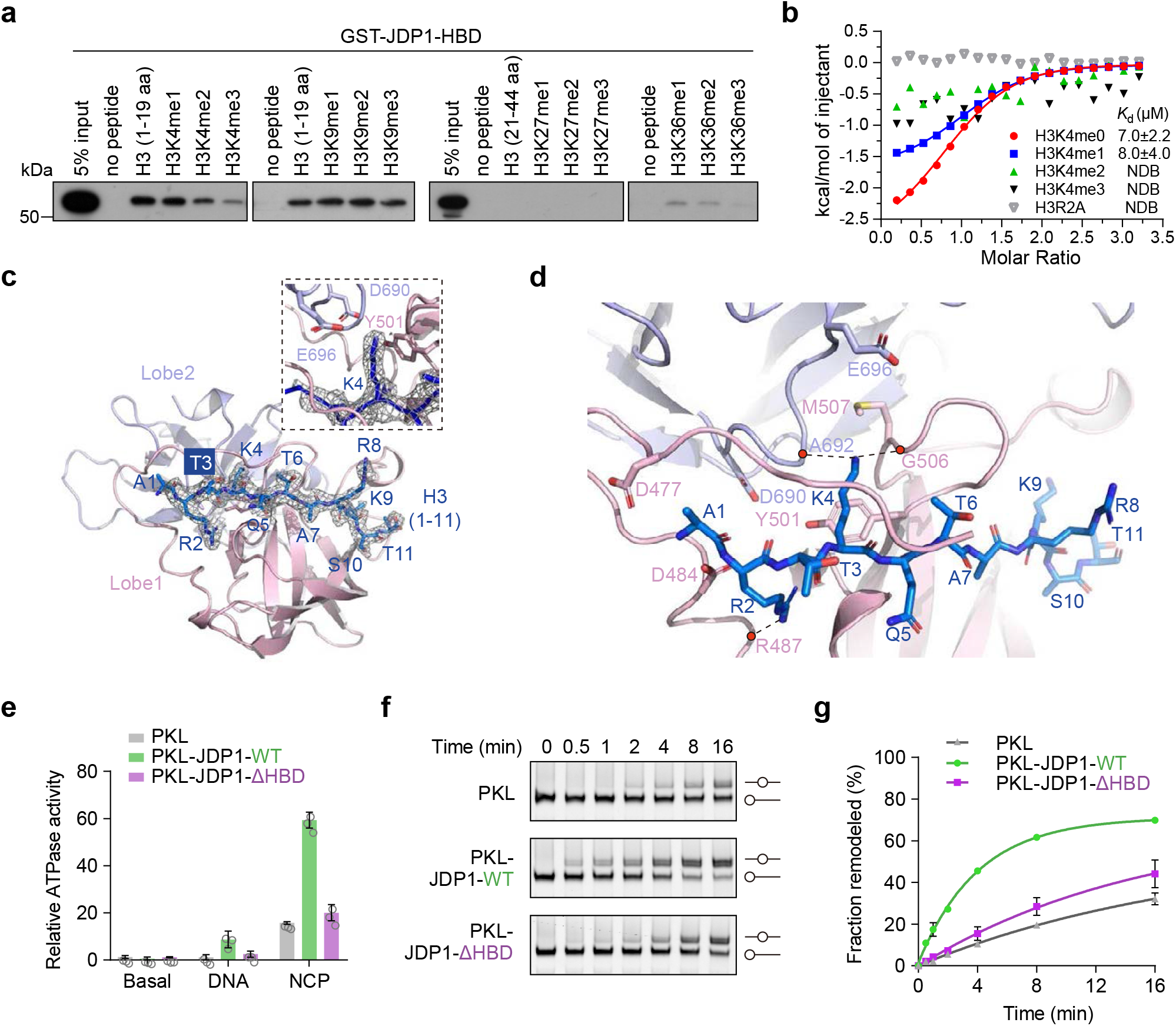
| JDP1 recognizes the histone H3 tail in an H3K4 hypomethylation-dependent manner via its histone-binding domain. (**a**) Pull-down assay showing the interaction between the JDP1-HBD (460-706 aa) and various histone H3 peptides. (**b**) ITC-based affinity measurement of the JDP1-HBD binding to H3K4me0, H3K4me1, H3K4me2, H3K4me3, and H3R2A peptides. NDB, no detectable binding. (**c**) Crystal structure of the JDP1-HBD in fusion with the H3 tail peptide, showing the electron density map (2*F*_o_-*F*_c_) around the H3 tail peptide at the contour level *σ* = 1. Local structure around H3K4 is shown on the top, with interacting residues highlighted in stick representation. (**d**) Local structure around the bound H3 peptide. The interacting residues are highlighted in stick representation. (**e**) Relative ATPase activities of PKL, PKL-JDP1-WT and PKL-JDP1-ΔHBD. Data are presented as mean ±SD (n = 3). (**f, g**) Chromatin-remodeling activities of PKL, PKL-JDP1-WT, PKL-JDP1-ΔHBD toward the 0N40 nucleosome. Representative gels are presented in (**f**), with quantification of the fractions of nucleosome slide shown in (**g**). Data are presented as mean ±SD (n = 3). The initial reaction rates (min^−1^) estimated by fitting and data are 0.03, 0.18 and 0.04 for PKL, PKL-JDP1-WT, PKL-JDP1-ΔHBD, respectively.

### Structural and functional analyses of the JDP1-HBD bound to the H3 tail

To elucidate the mechanism of H3 recognition, we first determined the crystal structure of HBD at a resolution of 2.1 Å, with R_free_/R_work_ refined to 23.1/26.8 (Extended Data Fig. 9a and Supplementary Table 2). The predicted structure matches the ground-true structure reasonably well, with a root-mean square deviation (RMSD) of 0.52 Å over 234 Cα atoms (Extended Data Fig. 9a). However, the H3-tail peptide could not be simply soaked into the crystals or co-crystallized with HBD. Analysis of the crystal lattice indicated that the predicted H3-binding pocket is blocked by the crystal contacts from the adjacent molecule (Extended Data Fig. 9b). To illustrate the specific mechanism of H3 recognition by HBD, we designed a construct with H3 (residues 1-19) in fusion with HBD (residues 469-706) based on the predicted H3-bound structure, and successfully determined the crystal structure of the fusion protein at a resolution of 2.2 Å, with R_free_/R_work_ refined to 20.1/23.4 (Fig. 5c, d and Supplementary Table 2). The structure shows that while HBD largely remains its folding upon H3 binding (Extended Data Fig. 9c), a highly acidic surface binds to the Ala1-Ser11 segment of the H3 tail (Fig. 5D and Extended Data Fig. 9d).

The structure reveals the detailed mechanism of H3 binding by HBD (Fig. 5c, d). The H3 tail is mainly bound by the first lobe of HBD (Lobe1), with H3K4 inserted into a pocket formed by the Lobe1-Lobe2 interface. The N-terminal amino group at the very end of H3 forms a hydrogen bond with Asp477 of Lobe1. Arg2 of H3 contributes markedly to the binding with its main chain interacting with the side chain of Asp484, and its side chain forming a hydrogen bond with the main chain of Arg487 in HBD (Fig. 5d and Extended Data Fig. 9e). Notably, the side chain of H3K4 packs against the hydrophobic side chains of Tyr501, and forms two hydrogen bonds with the main chains of Ala692 and Gly506 (Fig. 5d). Di- and tri-methylation of H3K4 are expected to disrupt these hydrogen bond interactions, providing a structural explanation for the observed sensitivity to H3K4 methylation (Fig. 5a, b). By contrast, the side chain of H3K9 is largely exposed to the solvent, consistent with the insensitivity to H3K9 methylation. Thus, the structural observation is fully consistent with the biochemical analyses and the ITC measurements above.

The H3K4 recognition mode by HBD is markedly different from those employed by canonical BAH domains, which typically engage methylated lysine residues through conserved aromatic cage structures (Extended Data Fig. 9f, g).^67^ The canonical aromatic cage residues are absent in HBD, with these positions substituted by residues L510 and S531 in Lobe1 and by P535, E617, and V637 in Lobe2 (Extended Data Fig. 9h, i). Instead, H3K4 binds to a newly formed pocket comprised of Lobe1 and Lobe2 in HBD (Fig. 5c, d).

To test the functional importance of HBD, we generated a HBD-truncated JDP1 variant and assessed its impact on the PKL-JDP1 complex activity *in vitro*. Deletion of the HBD caused a pronounced reduction in the ATPase and nucleosome remodeling activities (Fig. 5e-g). These results demonstrate that JDP1-mediated stimulation of PKL requires the histone-binding domain.

### The histone-binding domain is critical for compact chromatin with hypomethylated H3K4

Given that the HBD preferentially engage the histone H3 tail in an H3K4 hypomethylation-dependent manner, and that this interaction enhances PKL remodeling activity *in vitro* (Fig. 5), we hypothesized that PKL activity *in vivo* is modulated by H3K4 methylation status. To test this hypothesis, we classified PKL target loci into two groups based on changes in chromatin accessibility in the *jdp1/2* mutant. Group I loci exhibited increased chromatin accessibility in *jdp1/2* mutants relative to wild-type plants, whereas Group II loci showed no such increase. Notably, compared with Group II, Group I loci displayed significantly lower levels of the active histone mark H3K4me3, concomitantly with higher levels of the repressive mark H3K27me3 (Fig. 6a, b and Supplementary Data 6). These findings suggest that the chromatin-remodeling activity of the PKL-JDP complex is selectively promoted at genomic regions with low H3K4me3 levels to restrict chromatin accessibility.

**Fig. 6.**
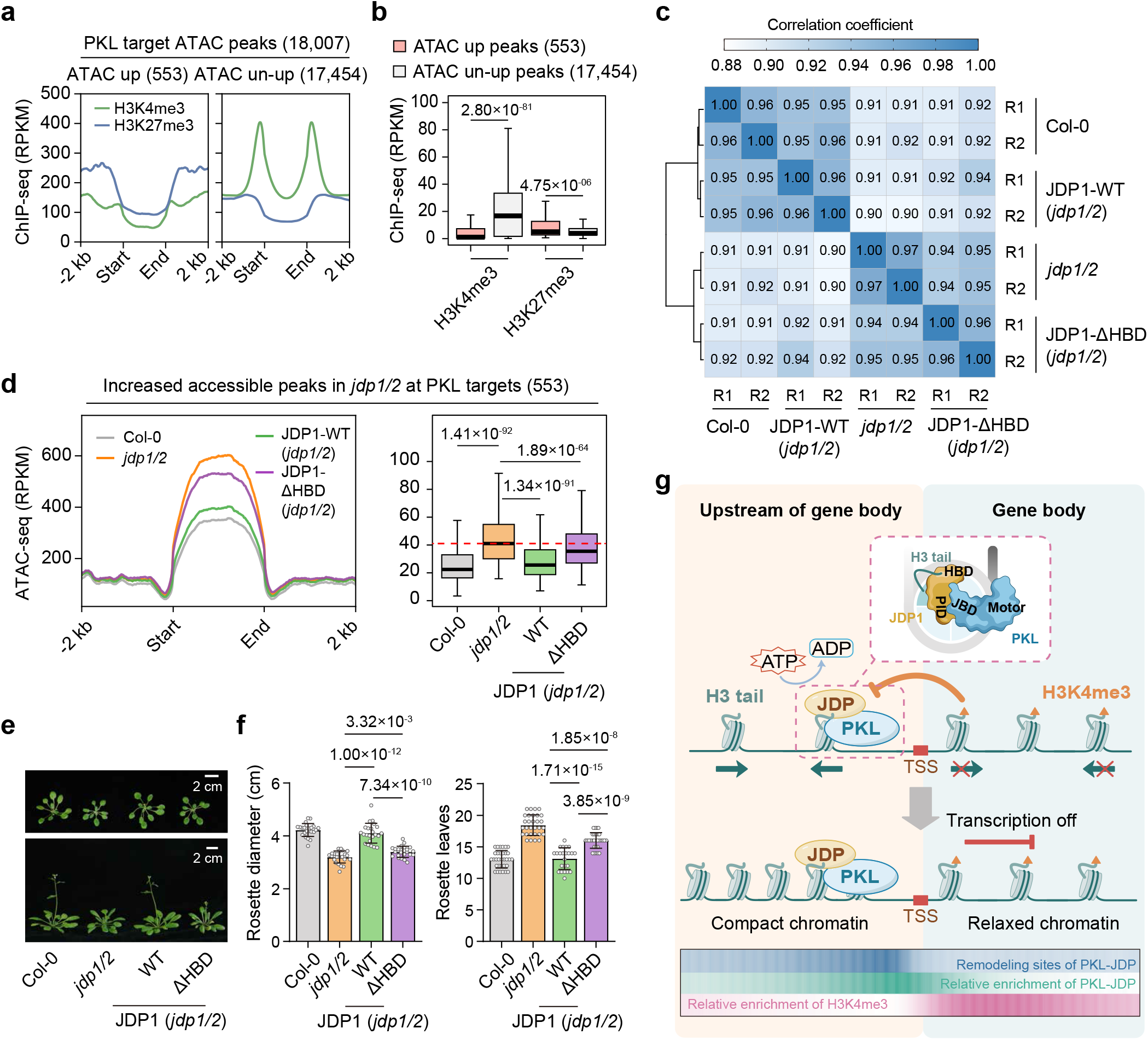
| Involvement of the HBD in JDP-dependent restriction of chromatin accessibility at hypomethylated H3K4 loci. (**a**) Metaplots depicting the distribution of H3K4me3 and H3K27me3 ChIP-seq signals at two categories of PKL target genomic loci. These loci were classified into two groups based on whether their ATAC-seq signals were upregulated in the *jdp1/2* mutant. (**b**) Boxplots depicting H3K4me3 and H3K27me3 levels at genomic intervals encompassing two subsets of ATAC-seq peaks (*jdp1/2*-upregulated and *jdp1/2*-un-upregulated) and their 500-bp flanking regions. *p* values were determined by a two-tailed Mann-Whitney U test (paired) for non-normally distributed data. Center lines and box edges are medians and the interquartile range (IQR), respectively. Whiskers extend within 1.5 times the IQR. (**c**) Heatmap showing pairwise correlation coefficients of all the ATAC-seq peaks among Col-0, *jdp1/2*, and complementation lines expressing JDP1-WT and JDP1-ΔHBD. The color bar shows the relationship between color shades and correlation values, with numerical coefficients shown in each heatmap cell. (**d**) Comparative analysis of chromatin accessibility in Col-0, *jdp1/2*, and JDP1-WT and JDP1-ΔHBD complementation lines. Analyses were performed using PKL target genomic loci with upregulated chromatin accessibility in the *jdp1/2* mutant relative to the wild-type Col-0 control. For boxplots, *p* values were determined using the two-tailed Mann-Whitney U test (paired) for non-normally distributed data. Center lines and box edges are medians and the interquartile range (IQR), respectively. Whiskers extend within 1.5 times the IQR. (**e, f**) Effects of HBD deletion on plant development and flowering time. Representative morphological phenotypes of 3-week-old plants (top) and 5-week-old plants (bottom) are shown in (**e**); quantification of rosette width (left) in 3-week-old plants and rosette leaf number (right) in 5-week-old plants is shown in (**f**). Data are presented as mean ±SD. *p* values were determined using a two-tailed Student’s t-test. Individual data points are represented by circles. (**g**) Working model of the PKL-JDP complex. The PKL-JDP complex associates with TSS-proximal regions. JDP binds the N-terminal tail of histone H3 lacking H3K4me3 and stimulates PKL-mediated chromatin compaction upstream of gene bodies. In contrast, H3K4me3 enrichment within gene bodies inhibits JDP binding to H3, thereby preventing PKL-mediated chromatin compaction. Color bars at the bottom indicate relative enrichment of PKL-JDP, H3K4me3, and PKL-JDP-dependent remodeling sites at TSS-proximal regions.

To dissect the contribution of the HBD to the chromatin-remodeling function of the PKL-JDP complex, we performed complementation analyses by generating transgenic lines expressing either *JDP1-WT* or *JDP1-ΔHBD* in the *jdp1/2* mutant background. Immunoblot assays revealed that the reduced PKL protein abundance observed in the *jdp1/2* double mutant was substantially rescued to comparable levels by both *JDP1-ΔHBD* and *JDP1-WT* transgenes (Extended Data Fig. 10a). These results indicate that JDP1-HBD is dispensable for maintaining PKL protein levels, supporting a model in which JDP1-HBD functions downstream of the PKL-JDP complex assembly.

Correlation analyses of ATAC-seq data revealed that chromatin accessibility in *JDP1-ΔHBD* transgenic plants more closely resembled that of the *jdp1/2* mutant than that of *JDP1-WT* transgenic plants or wild-type Col-0 controls (Fig. 6c and Extended Data Fig. 10b). Further analysis showed that expression of the *JDP1-WT* transgene reduced chromatin accessibility to levels comparable to wild type (Fig. 6d), indicating that JDP1 is required for restricting chromatin accessibility at these sites. The *JDP1-ΔHBD* transgene also reduced chromatin accessibility, but the effect was noticeably weaker than that of the *JDP1-WT* transgene (Fig. 6d and Extended Data Fig. 10c, d). Along the same line, while the *JDP1-WT* transgene fully rescued the reduced plant size and delayed flowering-time phenotypes of *jdp1/2* mutants, the *JDP1-ΔHBD* transgene conferred only partial complementation (Fig. 6e, f). These *in vivo* results are consistent with our *in vitro* nucleosome-centering assays (Fig. 5f, g), supporting the conclusion that HBD-mediated H3 interaction promotes the PKL remodeling activity.

## Discussion

This study reveals that a previously uncharacterized subfamily of J-domain proteins interacts with PKL to form a distinct PKL-JDP chromatin-remodeling complex. We demonstrate that these JDPs selectively recognize hypomethylated H3K4 tails, thereby stimulating PKL function to compact chromatin at distal promoters and intergenic regions that are depleted of the active histone mark H3K4me3 (Fig. 6g). Accumulating evidence indicates that arrays of neighboring nucleosomes enhance the activity of multiple chromatin-silencing factors, including heterochromatin protein HP1, the histone H3K27 methyltransferase complex PRC2, the DNA methyltransferase ZMET2, and the chromatin remodeler Isw1a.^68–70,42^ Our study implicates that the nucleosome arrays generated by the PKL-JDP complex create a chromatin environment permissive for H3K27 trimethylation and DNA methylation. This model is consistent with previous studies demonstrating that PKL contributes to PRC2-mediated H3K27me3 deposition and RNA-directed DNA methylation.^33,35^

Although H3K27me3 and DNA methylation are largely mutually exclusive repressive chromatin marks in Arabidopsis, both are characteristically depleted of the active histone modification H3K4me3.^36–38^ Recognition of hypomethylated H3K4 by JDPs therefore enables the PKL-JDP complex to selectively compact chromatin at regions lacking H3K4me3. PKL-mediated chromatin compaction subsequently facilitates the establishment of H3K27me3 and DNA methylation, linking H3K4 hypomethylation to two distinct repressive chromatin marks. Given that PKL regulates multiple developmental phase transitions by promoting H3K27 trimethylation and transcriptional repression at key developmental loci,^28,29^ the coupling of H3K4 hypomethylation to PKL-dependent chromatin remodeling mediated by JDPs provides a conceptual framework for understanding how H3K27me3 is initially established during diverse developmental phase transitions.

In mammals, CHD3-type chromatin remodelers contain N-terminal tandem PHD zinc fingers that recognize unmethylated H3K4 and methylated H3K9, both associated with repressive chromatin^71,72^. Although Arabidopsis PKL also harbors an N-terminal PHD zinc finger,^24^ our CL-MS analyses did not detect interactions involving the PHD zinc finger in the PKL-JDP-nucleosome complex. Consistent with this observation, deletion of the PHD zinc finger does not impair PKL function in Arabidopsis,^33,45^ arguing against an essential role for this domain. Instead, our structural analyses reveal that a conserved RYA motif located N-terminal to the PHD zinc finger engages the H2A-H2B surface of the nucleosome, inducing a subtle unwrapping of the exist DNA. Notably, the RYA motif is absent from metazoan CHD3-type chromatin remodelers but is broadly conserved in other chromatin remodelers, including Chd1, SMARCA4 and ISWI.^73–75^ Thus, although PKL is classified as a CHD3-type chromatin remodeler, it exhibits distinct structural and functional features that differentiate it from canonical metazoan CHD3-type enzymes.

CHD family chromatin remodelers generally induce regularly spaced nucleosome arrays.^76,14^ Whereas CHD1-type chromatin remodelers alone are sufficient for mediating the chromatin-remodeling activity, CHD3-type remodelers assemble into multi-subunit NuRD/Mi-2 complexes in metazoans.^15,17,77,20^ Although PKL belongs to the CHD3 family, it does not form a NuRD-like complex; instead, it associates with JDPs to assemble a distinct chromatin-remodeling complex. However, unlike NuRD, whose accessory subunits are largely dispensable for CHD catalytic activity,^1,78^ PKL activity is strongly enhanced by JDPs. These findings establish the PKL-JDP complex as functionally distinct from NuRD. Interestingly, the JDP-interacting region in PKL maps to a conserved region encompassing both the DUF1087 and SANT-like domains, which are conserved among CHD3-type remodelers across eukaryotes. Identification of this region as a JDP-interaction module therefore suggests a conserved regulatory function for this domain in CHD3-type remodelers beyond plants.

J-domain-containing proteins typically function as co-chaperones that stimulate the ATPase activity of HSP70 family proteins, thereby facilitating protein folding, assembly, and degradation.^79,80^ The Arabidopsis genome encodes more than 100 J-domain proteins, which are classified into four groups based on domain architecture.^81^ Several have been implicated in transcriptional regulation at methylated genomic regions.^82,83,58^ For example, the J-domain protein SILENZIO is recruited by the methyl-CpG-binding proteins MBD5 and MBD6 to mediate transcriptional silencing.^58^ Conversely, the J-domain proteins DNAJ1/2/3 (also known as SDJ1/2/3), form a complex with the DNA methylation readers SUVH1 and SUVH3 to antagonize transcriptional silencing at methylated loci.^82,83^ The JDPs identified in this study define a previously unrecognized subfamily of plant J-domain proteins characterized by an evolutionarily conserved histone-binding domain that is critical for enhancing PKL-dependent chromatin remodeling. In contrast to canonical J-domain proteins, whose J domains are essential for their HSP70-binding functions, disruption of the J domain in this subfamily does not impair their biological role in Arabidopsis. These findings indicate that the J domain in this subfamily functions through a mechanism distinct from classical HSP70-dependent pathways.

Importantly, we have uncovered a previously unrecognized HBD in the C-terminal region of JDPs that serves as a defining feature of PKL-interacting J-domain proteins. We demonstrate that the HBD specifically binds the N-terminal tail of histone H3 in its hypomethylated H3K4 state. Although the predicted structure of the HBD resembles that of the canonical histone-binding BAH domain, our crystal structure analysis reveals that the HBD bound to the histone H3 tail is fundamentally distinct from that of canonical BAH domains, which recognize H3K9me3 and H3K27me3 via aromatic cage motifs.^64–67^ The HBD is required to stimulate the nucleosome-remodeling activity of the PKL-JDP complex, providing a direct example of how histone modifications regulate chromatin remodeling. Given that the HBD is conserved in a subset of J-domain proteins across diverse plant lineages, including bryophytes, ferns, gymnosperms, and angiosperms, recognition of hypomethylated H3 tails by the HBD likely represents an evolutionarily conserved mechanism for regulating CHD3-mediated chromatin remodeling across a broad range of plant species.

## Materials and methods

### Plant materials and growth conditions

The *Arabidopsis* lines used in this study were in the Columbia-0 (Col-0) background unless otherwise indicated. The *pkl* mutant (CS339941) carries T-DNA insertion while *jdp1/2*, *jdp3*, *jdp4*, and *pap* mutations were generated by CRISPR-Cas9 technique. Higher-order mutants, including *jdp1/2/3*, *pkl/jdp1/2*, *pkl/jdp1/2/3*, *pkl/pap*, and *jdp1/2/3/pap* were generated by genetic crossing. The *jdp1/2/3/4* mutant was generated by CRISPR-Cas9-mediated disruption of *jdp4* in the *jdp1/2/3* triple-mutant background.

Transgenic plants expressing PKL, PKL-ΔRYA, JDP1, JDP1-A424D, JDP1-HPD/QQQ, JDP1-H/Q, JDP1-ΔHBD, JDP3, JDP4, or PAP proteins fused with FLAG or GFP epitope tags were generated in Col-0 or mutant background. Briefly, the modified pCAMBIA1305 vectors carrying the respective gene constructs were introduced into the *Agrobacterium tumefaciens* strain GV3101, and transgenic plants were obtained by Agrobacterium-mediated floral dip transformation of Col-0 or mutant plants. Genotypes and protein expression levels of transgenic lines were analyzed for verification.

The plants used for transcriptional silencing assays are of the ecotype C24. *Rd29A-LUC* was inserted into *ros1* mutant.^35^ The *ros1/pkl* was screened following EMS mutagenesis and *ros1/jdp1/2, ros1/jdp3, ros1/jdp1/2/3* were generated by CRISPR-Cas9 and genetic crossing. Primers for genotyping and transgene constructs are listed in Supplementary Data 7.

Plants were cultivated on Murashige and Skoog (MS) medium plates or in nutrient soil under long-day (16 h light/8 h dark) conditions at 22°C. For quantification of pickle roots, plants were grown on vertical MS medium plates under continuous illumination (24 h light) at 22°C.^23^

### Affinity purification with Mass spectrometry and co-immunoprecipitation assay

For FLAG-tagged protein affinity purification, 3 g of seedlings and inflorescences from transgenic plants were harvested after 11 days of growth and bolting. Tissues were frozen in liquid nitrogen and ground to a fine powder at 1000 rpm for 4 min. the powder was resuspended in plant lysis buffer (50 mM Tris-HCl [pH 7.5], 150 mM NaCl, 5 mM MgCl_2_, 10% glycerol, 0.1% NP-40, 0.5 mM DTT, 1 mM PMSF, protease inhibitor cocktail) and incubated with gentle rotation at 4°C for 2 h. Lysates were centrifuged at 18,000 × g and the supernatant was filtered through Miracloth (Millipore, 475855). 120 μL anti-FLAG M2 affinity gel beads (Sigma-Aldrich, A2220) were added into the filtrate and incubated at 4°C for 2.5 h with rotation.

Beads were collected by centrifugation at 800 ×g and washed six times with lysis buffer followed by eluted with 3 ×FLAG peptides (Sigma-Aldrich, F4799). Eluted proteins were resolved by 10% SDS-PAGE, and gels were fixed overnight in 50% ethanol and 10% acetic acid with gently agitation^84^. Silver staining was performed using the ProteoSilver Silver Stain Kit (PROT-SIL1, Sigma-Aldrich) according to the manufacturer’s instructions.

For mass spectrometry analysis, silver-stained gel bands were excised, destained, and digested with trypsin at 37°C overnight. Peptides were separated on a capillary column and analyzed by nano-electrospray ionization on a linear ion trap quadrupole mass spectrometer (Thermo Fisher Scientific). For mass spectrometry analysis, silver-stained gel bands were excised, destained, and digested with trypsin at 37°C overnight. Peptides were separated on a capillary column and analyzed by nano-electrospray ionization on a linear ion trap quadrupole mass spectrometer (Thermo Fisher Scientific). MS spectra were searched against the International Protein Index Arabidopsis protein database using Mascot. The protein-protein interactions were analyzed and visualized by GraphPad Prism 8.

For co-immunoprecipitation (co-IP) assay, 2 g of 11-day-old transgenic seedlings grown on MS medium plates were harvested. Seedlings were frozen in liquid nitrogen, ground to a fine powder, and resuspended in plant lysis buffer (as described for AP-MS). Lysates were centrifuged at 18,000 × g and the supernatant was filtered through Miracloth. An aliquot of 100 μL total proteins reserved as input, and the anti-FLAG M2 affinity gel beads were added to the remainder of the sample to capture FLAG-tagged proteins. Beads were collected by centrifugation, washed five times with lysis buffer, and eluted with 3 × Flag peptide (Sigma-Aldrich, F4799). Input and eluted samples were analyzed by immunoblotting using antibody of FLAG (Abclonal, AE005; 1:5000) or PKL (custom-made antibody from Huaan Biotechnology, 1:1000) as previously described.^26^

### Lipid staining

To assess the pickle root phenotype of *pkl* and *jdp1/2/3* mutants, lipid staining was performed using Sudan Red 7B (Sigma-Aldrich, 201618). 10-day-old seedlings were transferred in Sudan red staining solution (0.1% (w/v) Sudan Red 7B dissolved in PEG300 (Sigma-Aldrich, 202371) at 90°C for 1 h, followed by mixing with an equal volume 90% glycerol) for 3 h at room temperature, Seedlings were subsequently rinsed with 70% ethanol, as previously described.^33^ Seedlings displaying red-stained pickle roots were scored, and the percentage of affected seedlings was quantified to determine phenotype penetrance.

### Protein expression and purification

For *in vitro* protein expression and purification, full length coding sequences (CDS) of JDPs and PAP were amplified from cDNA and cloned into prokaryotic expression vectors (pGEX-6p-1 for GST-tagged proteins and pET-28a for His-tagged proteins). Truncated or mutated CDSs of *PKL* or *JDPs* were generated by multi-fragment ligation and homologous recombination approach. Recombinant GST- or His-tagged proteins were expressed in *Escherichia coli* strain T7 pLysY (DE3). X‘Bacteria cells were harvested, resuspended in lysis buffer (20 mM Tris-HCI [pH 7.5], 150 mM NaCl, 1 mM DTT), lysed by sonication, and subjected to affinity purification using glutathione Sepharose 4B beads (Cytiva, 17075601) for GST-tagged proteins or Ni-NTA His-bind Resin beads (EMD millipore corp, 70666) for His-tagged proteins. Purified proteins were analyzed by SDS-PAGE followed by Coomassie blue staining and immunoblotting.

For yeast expression, the full length CDS of *PKL* was cloned into pAT425 vector with a FLAG tag and expressed in *Saccharomyces cerevisiae* strain YPH499. Yeast cells were grown in synthetic defined medium lacking leucine (SD-L) at 30°C to an OD_600_ of approximately 1.0, harvested by centrifugation at 4,000 × g, and ground to a fine powder in liquid nitrogen. Cell lysates were prepared in yeast lysis buffer (50 mM Tris-HCl [pH 7.4]; 150 mM NaCl; 1 mM EDTA; 10% glycerol; 0.05% NP-40; 1 mM DTT; 1 mM PMSF; protease inhibitor cocktail) and clarified by centrifugation at 18,000 ×g.Anti-FLAG M2 affinity gel beads (Sigma-Aldrich, A2220) were added to the supernatant and incubated at 4°C with gentle rotation to capture recombinant PKL protein. Beads were washed five times with yeast lysis buffer then eluted with 3 ×Flag peptides (Sigma-Aldrich, F4799), as previously described.^85^

For obtaining co-expressed PKL-JDP complexes, the genes of *PKL* and *JDP* were sub-cloned into pCAG vectors, and transiently transfected into HEK293F cells with polyethyleneimines (PEI) (Yeasen, 40816ES01) for protein overexpression. To facilitate protein purification, His-tag and Strep-tag were added to C-termini of JDP1 and N-termini of PKL, respectively. The transfected cells were cultured in SMM 293-TII Expression Medium (Sino Biological Inc., M293TII) at 37°C supplemented with 5% CO_2_ in a shaker for 72 h. Cells were harvested, and then lysed by sonication in lysis buffer containing 50 mM Tris-HCl [pH 8.0], 500 mM NaCl, 10 mM imidazole, 2 mM MgCl_2_, 5% glycerol, 0.2% Tween 20 and protease inhibitor cocktail (5 μg/mL aprotinin, 2.5 μg/mL leupeptin and 2.5 μg/mL pepstatin). After centrifuged at 17,000 rpm at 4°C for 1 h, the supernatant was collected and applied to a HisTrap HP column (Cytiva, 29051021), washed by lysis buffer with 1 M NaCl and eluted with 250 mM imidazole. The elution was then loaded onto PreCap Streptactin (Smart-Lifesciences, SA053C11), washed by strep-wash buffer containing 50 mM Tris-HCl [pH 8.0], 300 mM NaCl and 5% glycerol, and eluted by strep-wash buffer supplemented by 8 mM D-desthiobiotin (Sigma-Aldrich, D1411). The elution sample was then applied to gel-filtration chromatography (Cytiva, Superdex-200) in buffer containing 20 mM Tris-HCl [pH 7.5], 150 mM NaCl and 2 mM DTT. PKL-JDP containing fractions were collected and concentrated to around 1.5 mg/mL. Primers used for protein expression constructs are listed in Supplementary Data 7.

### Yeast two-hybrid assay

Yeast two-hybrid (Y2H) assay were performed using modified pGBKT7 and pGADT7 vectors harboring full-length or truncated CDS of *PKL*, *JDPs* and *PAP*. Plasmids were propagated in *Escherichia coli* strain DH5α were introduced into yeast strains Y187 and AH109, respectively. Positive Y187 transformants were selected on synthetic dropout medium lacking tryptophan (SD-W), and positive AH109 transformants were selected on synthetic dropout medium lacking leucine (SD-L). Primers used for Y2H constructs are listed in Supplementary Data 7. AH109 and Y187 transformants were mated in yeast peptone dextrose adenine (YPDA) liquid medium overnight. Diploid yeast cells were plated on synthetic dropout medium lacking leucine and tryptophan (SD-W-L) to assess mating efficiency and on synthetic dropout medium lacking leucine, tryptophan, and histidine (SD-W-L-H) to evaluate protein-protein interaction, as previously described.^86^

### *In vitro* pull-down assay

For *in vitro* pull-down assays, purified FLAG-, GST-, and/or His-tagged proteins were mixed in tubes in reaction buffer (20 mM Tris-HCl [pH 7.5]; 100 mM NaCl; 1 mM DTT) and incubated with gentle rotation at 4°C for 4 h. Prior to the addition of affinity added, 10% of incubation sample was reserved as input control. The remaining samples were incubated with anti-FLAG M2 affinity gel beads or glutathione Sepharose beads for 1 h at 4°C. Beads were collected by centrifugation, washed five times with reaction buffer, and eluted with 3×FLAG peptide (Sigma-Aldrich, F4799) or 20 mM L-glutathione reduced (Sigma-Aldrich, G4251) for 30 min at 4°C. Input and eluted samples were analyzed by SDS-PAGE followed by Coomassie Brilliant Blue staining or immunoblotting using anti-FLAG (Abclonal, AE005; 1:5,000), anti-GST (Abmart, M20007L; 1:5,000), anti-His (Origene, F013; 1:5,000) or anti-PKL (custom-made antibody produced by Huaan Biotechnology; 1:1,000) antibodies.

### RNA extraction and RNA-seq analysis

For RNA extraction of seedlings and rosettes, 0.1 g of 7-, 11-day-old seedlings, and 12 plants of 21-day-rosettes from Col-0, *pkl*, *jdp1/2/3*, *jdp1/2/3/4* (11-day-old only for *jdp1/2/3/4*) were harvested. Samples were quickly frozen in liquid nitrogen, ground to a fine powder, and lysed in 1 mL TRIzol Reagent (Invitrogen, 15596018CN). Lysates were centrifuged at 12,000 ×g, and RNA in the aqueous phase was precipitated with an equal volume of isopropanol at -20°C for 30 min. The precipitate was collected by centrifugation and washed by 80% ethanol. RNA samples were sent to Novogene for library generation and sequencing using an Illumina NovaSeq 6000 platform (paired-end, 150bp). At least three independent biological replicates were performed. For RNA extraction of imbibed seeds and 3-day-old plants, 0.05 g of Col-0, *pkl*, *jdp1/2/3* were harvested. The RNA was extracted using the plant seed ultra-speed RNA extraction kit (Mei5bio, MF737). The experimental procedure was performed in accordance with the manufacturer’s instructions. In brief, samples were ground into a fine powder in liquid nitrogen and resuspended in buffer S1 buffer supplemented with β-mercaptoethanol. After mixing, samples were centrifuged at 13,000 × g for 10 min at 4°C, and ∼600 μL supernatant was collected. Buffer S2 and chloroform were added, mixed, incubated for 5 min, and centrifuged. The aqueous phase was transferred and mixed with 300 μL isopropanol, then loaded onto a spin column. The column was washed twice with WB buffer, and RNA was eluted with 50 μL EB buffer.

For RNA-seq analysis, adaptors and low-quality reads were removed, and clean reads were aligned to *Arabidopsis thaliana* genome (TAIR 10) using HISAT2 (v2.1.0).^87^ DEGs were identified using the R (v4.3.0) package edgeR (v3.42.4) with |log_2_FC| > 0.5 and FDR < 0.05.^88^ The GO enrichment analysis was performed using the GO enrichment function in the R package clusterProfiler (v4.8.1). Venn diagrams were generated using ggvenn (v0.1.10). Heatmaps of DEGs were generated using the R package gplots (v3.1.3.1). The correlation heatmap, bar plot, scatter plots and bubble charts were generated using the R package ggplot2 (v3.5.0).

### Chromatin immunoprecipitation (ChIP) assay and ChIP-seq analysis

For ChIP assay, 4 g of 11-day-old seedlings per replicate were harvested from MS medium plates. GFP-tagged transgenic plants in mutant backgrounds were used for protein target analysis. The experiment procedures were performed as previously described.^89^ Seedlings were cross-linked with 1% formaldehyde under vacuum quenched with 0.125 M glycine was added to terminate. Nuclei were isolated by grinding and sequential extraction in in extraction buffer I (10 mM Tris-HCl [pH 8.0], 10 mM MgCl_2_, 0.4 M sucrose, 0.25% Triton X-100, 1 mM DTT, 0.1 mM PMSF, protease inhibitor cocktail), washed in extraction buffer II (10 mM Tris-HCl [pH 8.0], 10 mM MgCl_2_, 0.25 M sucrose, 1% Triton X-100, 0.1 mM DTT, 0.1 mM PMSF, 1% protease inhibitor cocktail). and layered on buffer extraction III (10 mM Tris-HCl [pH 8.0], 2 mM MgCl_2_, 1.7 M sucrose, 0.15% Triton X-100, 1 mM DTT, 0.1 mM PMSF, protease inhibitor cocktail) for centrifugation at 13,000 ×g for 1 h.

Nuclei were sonicated in a 1:2 mixture of lysis buffer (50 mM Tris-HCl [pH 8.0], 10 mM EDTA, 0.1 mM DTT, 0.1 mM PMSF, protease inhibitor cocktail) and dilution buffer (16.7 mM Tris-HCl [pH 8.0], 1.2 mM EDTA, 167 mM NaCl, 1.1% Triton X-100, 1 mM DTT, 0.1 mM PMSF, protease inhibitor cocktail) for 23-28 cycles (30s on/30 s off) using a Bioruptor sonicator and clarified by centrifugation at 13,000 g for 15 min. The supernatant was diluted to <0.01% SDS and incubated with antibodies of GFP (Abcam, Ab290) for 12 h at 4°C followed by Dynabeads of protein A (ThermoFisher, 10001D) at 4°C for 4 h. The beads-antibody-chromatin fragments combination products were washed sequentially with low salt wash buffer (20 mM Tris-HCl [pH 8.0], 2 mM EDTA, 150 mM NaCl, 0.1% SDS, 1% Triton X-100), high salt wash buffer (20 mM Tris-HCl [pH 8.0], 2 mM EDTA, 500 mM NaCl, 0.1% SDS, 1% Triton X-100), LiCl wash buffer (10 mM Tris-HCl [pH 8.0], 1 mM EDTA, 0.25 M LiCl, 1% sodium deoxycholate, 1% NP-40) and TE wash buffer (10 mM Tris-HCl [pH 8.0], 1 mM EDTA [pH 8.0]). Chromatin fragments were eluted with 100 mM NaHCO3 containing 1% SDS and reverse cross-linked with 200 mM NaCl at 65°C overnight. DNA was extracted using phenol:chloroform:isoamyl alcohol (25:24:1) and ethanol precipitated overnight at -20°C. DNA libraries were prepared using the VAHTS Universal DNA Library Prep Kit for Illumina V4 (Vazyme, ND610) and sent to Novogene for high-throughput sequencing on the Illumina NovaSeq platform.

For ChIP-seq analysis, the clean data were mapped to the *Arabidopsis thaliana* genome (TAIR10) by Bowtie2 (v2.3.4) with one mismatch allowed (Langmead & Salzberg, 2012). PCR duplicates were removed with Picard Tools (v2.23.0). GFP ChIP-seq peaks were called with MACS2 (v2.2.7.1) in R.^90^ H3K4me3 and H3K27me3 peaks were identified using SICER2 (v1.0.2).^91^ At least two independent biological replicates were performed. Read counts were normalized reads per kilobase per million mapped reads (RPKM) by the number of clean reads mapped to the genome in each library. Venn diagrams were generated using ggvenn (v0.1.10). Heatmaps and metaplots were generated using DeepTools (v3.5.1). The bar plots and boxplots were drawn using the R package ggplot2 (v3.5.0).

### ATAC-seq and analysis

For ATAC-seq, ∼0.1 g plants of 11-day-old seedlings per replicate were harvested from MS medium plates. Seedlings were finely chopped on ice in pre-cooled nuclei extraction buffer (10 mM Tris-HCl [pH 8.0], 0.25 M sucrose, 10 mM MgCl2, 0.3% Triton X-100 and protease inhibitor cocktail). The homogenate was filtered through a 40 μm cell strainers and centrifuged at 1,000 ×g for 10 min at 4°C to isolate nuclei. The collected nuclei were washed and resuspended in nuclei purification buffer (20 mM MOPS [pH 7.0], 40 mM NaCl, 90 mM KCl, 2 mM EDTA, 0.5 mM EGTA, 0.5 mM spermidine, 0.2 mM spermine and protease inhibitor cocktail). Nuclei were stained with DAPI and counted using a hemocytometer under an optical microscope, and approximately 10,000-50,000 nuclei were used for subsequent Tn5 transposition at 37°C with intermittent mixing for 30 min. Following DNA purification, ATAC-seq libraries were prepared using the TruePrep DNA Library Prep Kit V2 for Illumina (Vazyme, TD501) and subjected to high-throughput sequencing by Novogene, as previously described.^92^

For ATAC-seq analysis, clean reads generated by Novagene were trimmed to remove adapter sequences and low-quality reads, and then aligned to the Arabidopsis reference genome (TAIR10) using Bowtie2 (v2.3.4) with up to one mismatch allowed.^93^ Probable PCR duplicates were removed using MarkDuplicates of Picard Tools (v2.23.0). The peaks were identified using MACS2 (v2.2.7.1) with the following parameters: -f BAMPE -keep-dup all.^90^ At least two independent biological replicates were performed in this study. Differentially accessible genomic regions between wild type and mutants were identified using the DiffBind package (v3.10.1) in R^94^. Venn diagrams, metaplots, and MA plots were generated using ggvenn (v0.1.10), DeepTools (v3.5.1) and DiffBind (v3.10.1), respectively. The bar plots, scatter plots, boxplots, volcano plots and correlation heatmap were drawn using the R package ggplot2 (v3.5.0).

### *In vitro* peptide binding assay

For the peptide-protein binding assays, 1 μg biotinylated histone peptides, including unmodified H3 (1-19 aa), H3K4me1 (1-21 aa), H3K4me2 (1-21 aa), H3K4me3 (1-21 aa), H3K9me1 (1-21 aa), H3K9me2 (1-21 aa), H3K9me3 (1-21 aa), unmodified H3 (21-44 aa), H3K27me1 (21-44 aa), H3K27me2 (21-44 aa), H3K27me3 (21-44 aa), H3K36me1 (21-44 aa), H3K36me2 (21-44 aa) and H3K36me3 (21-44 aa) (SciLight Peptide), were incubated with 5 μg purified GST-JDP1-HBD proteins were incubated in 300 μL pre-cold binding buffer (50 mM Tris-HCl [pH 7.5], 150 mM NaCl, 0.05% NP - 40) for 4 h. An aliquot of each reaction was reserved as input prior to the addition of 20 μL Streptavidin MagneSphere Paramagnetic Particles (Promega, Z5481). The mixtures were gently rotated for1 h at 4°C, after which the beads were collected using a magnet and washed three to four times with binding buffer. Samples were boiled at 98°C for 10 min in 40 μL 1× SDS loading buffer, as previously described.^95^ All the samples were analyzed by immunoblotting using an antibody of GST (Abmart, M20007L; 1:5,000).

### Alphafold3 prediction

AlphaFold3 was used to predict the interaction between PKL and JDP1. The predicted models were evaluated using standard confidence metrics, including the inter-chain predicted aligned error (PAE), per-residue local distance difference test (pLDDT), and interface template modeling score (ipTM), which collectively reflect the accuracy of intermolecular docking and local structural quality. High-confidence intermolecular binding was defined based on widely adopted thresholds: inter-chain PAE values below 10 Å for reliable domain–domain docking and interface residue pLDDT scores above 70 for robust side-chain conformation prediction.

### ITC measurements

The interaction of HBD with unmodified or modified H3 peptides (H3 (1-19), H3 (1-21) K4me1, H3 (1-21) K4me2, H3 (1-21) K4me3; synthesized by SciLight Peptide) in this paper were quantitatively assessed by ITC assay. The solvent of purified proteins was exchanged into ITC buffer (20 mM Tris-HCl [pH 7.5] and 150 mM NaCl, filtered by 0.22 μm filter) through Amicon Ultra Centrifugal Filters (Millipore, UFC901096) at 4 °C. The concentrations of the H3 peptides (1,000 μM) in the syringe were about 17-fold higher than the concentration of HBD in the cell (60 μM). ITC reactions were subjected by a standard approach and the data visualised by GraphPad Prism 8.

### Nucleosome reconstitution

Histone octamer was assembled from purified histones as described.^96^ The ‘601’ DNA fragment with 40 bp linker DNA at one end was used to assemble the nucleosomes (0N40). Similarly, the nucleosome reconstituted with the ‘601’ DNA with flanking linker DNA 20 bp and 40 bp at either end (20N40) was used for structure determination. Sequences used for linker DNA are listed in Supplementary Data 7.

### Nucleosome centering and ATPase activities

Nucleosome centering activities were performed as described previously.^97^ Reactions were performed with 20 nM Cy5-labeled 0N40 nucleosome and 20 nM proteins in the buffer (50 mM KCl, 20 mM Tris-HCl [pH 7.5], 5 mM MgCl_2_, 0.1 mg/mL bovine serum albumin, 5% glycerol, 1 mM DTT) with 2 mM ATP. The reactions were stopped at the indicated time points with 20 mM Tris-HCl [pH 7.5], 2mM EDTA and 800 ng/μL sperm DNA. Fractions were run on 8% native TBE polyacrylamide gels in 0.25 × TBE for 180 min at 150 V at 4 °C. Gels were imaged using an Amershan Typhoon 5 variable mode imager (Cytiva, 29187191). Band intensities were quantified in Quantity One software. The fraction of nucleosomes centered at a given time point was determined by the ratio of remodeled nucleosomes to total nucleosome intensity in the same lane of the gels. The reaction rate constants were fitted to a single exponential decay using Origin v.9.2. Measurement of ATP hydrolysis was performed as previously described, using proteins (60 nM) and double-stranded DNA (300 nM).

### CL-MS analyses

The CL-MS analyses were performed as described previously.^98^ Purified PKL-JDP complex (3.2 μM) was preincubated with the 20N40 nucleosome (1.5 μM) for 4 h. The PKL-JDP-Nucleosome Core Particle (NCP) complex was cross-linked with 5 mM bis(sulfosuccinimidyl)suberate (BS3) at room temperature for 30 min, which was then quenched with 100 mM Tris-HCl [pH 8.0]. Sample processing and data analyses were done as described.

### Electron microscopy sample preparation and data collection

The PKL-JDP-NCP complex was obtained by mixing 3.8 μM proteins with 1.9 μM 20N40 in the presence of ADP-BeF_x_. The complex was purified and stabilized using the GraFix protocol similarly as described before.^99^

For cryo-EM grid preparation, Quantifoil gold R2/1 grids with 200 mesh size were glow discharged in air for 30 s using a PDC-32G-2 Plasma Cleaner set to a low power. Samples (4 μL) were blotted for 3.0 s at −2 force before being plunge-frozen in liquid ethane with a FEI Vitrobot IV at 8 °C and 100% humidity. Grids were examined and screened using an FEI Tecnai Arctica operated at 200 kV. Cryo-EM data were collected using an FEI Titan Krios operated at 300 kV with a K3 direct electron detector (Gatan, 1025), at a magnification of ×29,000 for a final pixel size of 0.485 Å/pixel with the defocus values ranging from −1.2 to −1.8 μm. The total electron dose was 50 e−/Å^2^ fractionated in 32 frames.

### Image processing and model building

A total of 5,475 dose-fractionated image stacks were firstly aligned using MotionCo2 with twofold binning, and the contrast transfer function parameters were estimated using CTFFIND4. Particle picking, two-dimensional (2D) classification and three-dimensional (3D) classification were carried out in CryoSPARC v4. The initial picked particles were extracted with fourfold binning to increase signal to noise ratio. After multiple rounds of 2D classification, a total of 795,110 particles were selected and subjected to further processing. Heterogeneous refinement was performed using reference maps generated from ab-initio reconstruction, junk particles without well-defined nucleosome were removed at these steps. A set of 382,204 particles were then re-extracted without binning and subjected to CTF refinement, yielding a structure of PKL-NCP at an overall resolution of 3.1 Å. Some weak density appeared at a low contour level on the H2A-H2B acidic patch of the nucleosome. The particles were further classified with spherical mask focused on RYA, which was reconstructed to a resolution of 3.0 Å. The initial model was built by fitting the maps in Chimera using the predicted structures by AlphaFold3. The atomic models for the rest of the molecules were built manually in Coot. The structures were refined using Phenix with secondary structure constrains.

### Crystallization, data collection and structure solution

Crystals of HBD were grown at 4 °C by hanging drop vapor diffusion above a reservoir solution of 0.2 M MgCl_2_, 0.1 M Tris-HCl [pH 8.5], 25% (w/v) PEG3350, with protein to reservoir volume ratio 1:1. Crystals were harvested in cryoprotectant containing 10% glycerol and then flash frozen in liquid nitrogen. Crystals of H3-HBD were incubated with 0.2 M potassium sodium tartrate tetrahydrate and 20% (w/v) PEG3350 using sitting drop vapor diffusion at 4 °C. Crystals were harvested in reservoir buffer with 15% glycerol as cryo-protectant and then flash-frozen in liquid nitrogen.

Diffraction data from crystals of HBD and HBD-H3 were collected at the Shanghai Synchrotron Radiation Facility (SSRF), beamline BL02U1 and BL19U1, and processed with the autoPROC and autoPX package, respectively. The structures were solved by molecular replacement using the predicted structure of HBD as the initial searching model. The rest of the models were built manually with Coot. Refinement was performed with Phenix. The final structure of HBD domain was refined to 2.12 Å, with R_work_/R_free_=0.231/0.268, Ramachandran outlier 0.00%, allowed 4.76%, and favored 95.24%. The final structure of H3-HBD was refined to 2.24 Å, with R_work_/R_free_ = 0.201/0.234, Ramachandran outlier 0.00%, allowed 4.64%, and favored 95.36%.

## Data, code, and materials availability

Raw ChIP-seq, RNA-seq and ATAC-seq data are available from the Genome Sequence Archive database in the National Genomics Data Center (http://ngdc.cncb.ac.cn/) under accession number CRA032820 (https://ngdc.cncb.ac.cn/gsa/s/dt1d07tm). The same datasets are also available from the GEO database under accession codes GSE308893 (ChIP-seq; secure token: uzytmswaprslhix), GSE308894 (RNA-seq; secure token: ohwxgcgotneljqn), and GSE308895 (ATAC-seq; secure token: spenaciktdcxlmz). H3K4me3 ChIP-seq data in Col-0 is from GEO database under the accession code GSE281399. The AlphaFold3 predicted models of PKL-JDP1 and PBD-PID are available in ModelArchive (https://modelarchive.org) with the accession codes ma-4ouua and ma-sjf2u, respectively. Coordinates and EM maps are available from the Electron Microscopy Data Bank and Protein Data Bank under accession numbers EMD-68253 and 22FZ (PKL-NCP), EMD-83227 (PKL-motor), EMD-68244 (RYA-NCP), 22FI (HBD) and 22EZ (HBD-H3).

The Arabidopsis sequence information used in this study was derived from TAIR10. The accession numbers of genes reported in this study are AT2G25170 (*PKL*), AT2G05250 (*JDP1*), AT2G05230 (*JDP2*), AT2G25560 (*JDP3*), AT5G53150 (*JDP4*), AT1G01730 (*PAP*), AT1G08600 (*ATRX*) AT5G61380 (*TOC1*), AT5G37260 (*CIR1*), AT5G60100 (*PRR3*), AT3G46640 (*LUX*), AT1G25560 (*TEM1*), AT3G07650 (*COL9*), AT4G16250 (*PHYD*), AT3G54990 (*SMZ*), AT4G36920 (*AP2*), AT3G20810 (*JMJ30*), AT1G66050 (*VIM2*), AT1G47990 (*GA2OX4*), AT5G17490 (*RGL3*), AT5G64170 (*LNK1*), AT3G54500 (*LNK2*), AT5G02810 (*PRR7*), AT2G46790 (*PRR9*), AT5G60910 (*AGL8*), AT4G24540 (*AGL24*), AT4G02780 (*GA1*), AT1G15550 (*GA3OX1*), AT3G15270 (*SPL5*), AT2G42200 (*SPL9*), AT5G04275 (*MIR172B*), AT2G43010 (*PIF4*). All materials are available upon reasonable request.

## Acknowledgements

This work was supported by the National Natural Science Foundation of China to Xin-Jian He (grant number: 32025003 and 32470628).

## Author contributions

X.-J.H and Z.C. conceived the project and design the experiments. B.-B.G., Z.Y., P.D., Y.S., Y.-X.L., J.-X.L., X.-M.S., J.G., Z.-Z.L., X.-Y.L., L.L., and S.C. performed the experiments and data analysis. D.-Y.Y. and Y.-N.S. performed bioinformatics analysis; B.-B.G., Z.Y., P.D., Z.C., and X.-J.H. analyzed the data and wrote the manuscript. Competing Interests: The authors declare no competing interests.

## References

1. Eustermann, S., Patel, A.B., Hopfner, K.-P., He, Y., and Korber, P. (2024). Energy-driven genome regulation by ATP-dependent chromatin remodellers. Nat Rev Mol Cell Biol 25, 309–332. 10.1038/s41580-023-00683-y.

2. Thurman, R.E., Rynes, E., Humbert, R., Vierstra, J., Maurano, M.T., Haugen, E., Sheffield, N.C., Stergachis, A.B., Wang, H., Vernot, B., et al. (2012). The accessible chromatin landscape of the human genome. Nature 489, 75–82. 10.1038/nature11232.

3. Kharchenko, P.V., Alekseyenko, A.A., Schwartz, Y.B., Minoda, A., Riddle, N.C., Ernst, J., Sabo, P.J., Larschan, E., Gorchakov, A.A., Gu, T., et al. (2011). Comprehensive analysis of the chromatin landscape in Drosophila melanogaster. Nature 471, 480–485. 10.1038/nature09725.

4. Zhong, Z., Feng, S., Duttke, S.H., Potok, M.E., Zhang, Y., Gallego-Bartolomé, J., Liu, W., and Jacobsen, S.E. (2021). DNA methylation-linked chromatin accessibility affects genomic architecture in *Arabidopsis*. Proc. Natl. Acad. Sci. U.S.A. 118, e2023347118. 10.1073/pnas.2023347118.

5. Zhang, W., Zhang, T., Wu, Y., and Jiang, J. (2012). Genome-Wide Identification of Regulatory DNA Elements and Protein-Binding Footprints Using Signatures of Open Chromatin in *Arabidopsis*. Plant Cell 24, 2719–2731. 10.1105/tpc.112.098061.

6. Klemm, S.L., Shipony, Z., and Greenleaf, W.J. (2019). Chromatin accessibility and the regulatory epigenome. Nat Rev Genet 20, 207–220. 10.1038/s41576-018-0089-8.

7. Chen, Y., Liang, R., Li, Y., Jiang, L., Ma, D., Luo, Q., and Song, G. (2024). Chromatin accessibility: biological functions, molecular mechanisms and therapeutic application. Sig Transduct Target Ther 9, 340. 10.1038/s41392-024-02030-9.

8. Niederhuber, M.J., and McKay, D.J. (2021). Mechanisms underlying the control of dynamic regulatory element activity and chromatin accessibility during metamorphosis. Current Opinion in Insect Science 43, 21–28. 10.1016/j.cois.2020.08.007.

9. Wu, J., Huang, B., Chen, H., Yin, Q., Liu, Y., Xiang, Y., Zhang, B., Liu, B., Wang, Q., Xia, W., et al. (2016). The landscape of accessible chromatin in mammalian preimplantation embryos. Nature 534, 652–657. 10.1038/nature18606.

10. Tian, H., Li, Y., Wang, C., Xu, X., Zhang, Y., Zeb, Q., Zicola, J., Fu, Y., Turck, F., Li, L., et al. (2021). Photoperiod-responsive changes in chromatin accessibility in phloem companion and epidermis cells of Arabidopsis leaves. The Plant Cell 33, 475–491. 10.1093/plcell/koaa043.

11. Guo, J., Liu, Z.-Z., Su, X.-M., Su, Y.-N., and He, X.-J. (2025). The SAS chromatin-remodeling complex mediates inflorescence-specific chromatin accessibility for transcription factor binding. Nucleic Acids Research 53, gkaf316. 10.1093/nar/gkaf316.

12. Khorasanizadeh, S. (2004). The nucleosome: from genomic organization to genomic regulation. Cell 116, 259–272. 10.1016/s0092-8674(04)00044-3.

13. Luger, K., Dechassa, M.L., and Tremethick, D.J. (2012). New insights into nucleosome and chromatin structure: an ordered state or a disordered affair? Nat Rev Mol Cell Biol 13, 436–447. 10.1038/nrm3382.

14. Clapier, C.R., Iwasa, J., Cairns, B.R., and Peterson, C.L. (2017). Mechanisms of action and regulation of ATP-dependent chromatin-remodelling complexes. Nat Rev Mol Cell Biol 18, 407–422. 10.1038/nrm.2017.26.

15. Tong, J.K., Hassig, C.A., Schnitzler, G.R., Kingston, R.E., and Schreiber, S.L. (1998). Chromatin deacetylation by an ATP-dependent nucleosome remodelling complex. Nature 395, 917–921. 10.1038/27699.

16. Zhang, Y., Ng, H.-H., Erdjument-Bromage, H., Tempst, P., Bird, A., and Reinberg, D. (1999). Analysis of the NuRD subunits reveals a histone deacetylase core complex and a connection with DNA methylation. Genes & Development 13, 1924–1935. 10.1101/gad.13.15.1924.

17. Xue, Y., Wong, J., Moreno, G.T., Young, M.K., Côté, J., and Wang, W. (1998). NURD, a Novel Complex with Both ATP-Dependent Chromatin-Remodeling and Histone Deacetylase Activities. Molecular Cell 2, 851–861. 10.1016/S1097-2765(00)80299-3.

18. Yoshida, T., Hazan, I., Zhang, J., Ng, S.Y., Naito, T., Snippert, H.J., Heller, E.J., Qi, X., Lawton, L.N., Williams, C.J., et al. (2008). The role of the chromatin remodeler Mi-2β in hematopoietic stem cell self-renewal and multilineage differentiation. Genes Dev. 22, 1174–1189. 10.1101/gad.1642808.

19. Hoffmeister, H., Fuchs, A., Erdel, F., Pinz, S., Gröbner-Ferreira, R., Bruckmann, A., Deutzmann, R., Schwartz, U., Maldonado, R., Huber, C., et al. (2017). CHD3 and CHD4 form distinct NuRD complexes with different yet overlapping functionality. Nucleic Acids Research 45, 10534–10554. 10.1093/nar/gkx711.

20. Wade, P.A., Gegonne, A., Jones, P.L., Ballestar, E., Aubry, F., and Wolffe, A.P. (1999). Mi-2 complex couples DNA methylation to chromatin remodelling and histone deacetylation. Nat Genet 23, 62–66. 10.1038/12664.

21. Fujita, N., Jaye, D.L., Geigerman, C., Akyildiz, A., Mooney, M.R., Boss, J.M., and Wade, P.A. (2004). MTA3 and the Mi-2/NuRD Complex Regulate Cell Fate during B Lymphocyte Differentiation. Cell 119, 75–86. 10.1016/j.cell.2004.09.014.

22. Lai, A.Y., and Wade, P.A. (2011). Cancer biology and NuRD: a multifaceted chromatin remodelling complex. Nat Rev Cancer 11, 588–596. 10.1038/nrc3091.

23. Ogas, J., Cheng, J.-C., Sung, Z.R., and Somerville, C. (1997). Cellular Differentiation Regulated by Gibberellin in the *Arabidopsis thaliana pickle* Mutant. Science 277, 91–94. 10.1126/science.277.5322.91.

24. Ogas, J., Kaufmann, S., Henderson, J., and Somerville, C. (1999). PICKLE is a CHD3 chromatin-remodeling factor that regulates the transition from embryonic to vegetative development in *Arabidopsis*. Proc. Natl. Acad. Sci. U.S.A. 96, 13839–13844. 10.1073/pnas.96.24.13839.

25. Aichinger, E., Villar, C.B.R., Di Mambro, R., Sabatini, S., and Köhler, C. (2011). The CHD3 Chromatin Remodeler PICKLE and Polycomb Group Proteins Antagonistically Regulate Meristem Activity in the *Arabidopsis* Root. The Plant Cell 23, 1047–1060. 10.1105/tpc.111.083352.

26. Jing, Y., Zhang, D., Wang, X., Tang, W., Wang, W., Huai, J., Xu, G., Chen, D., Li, Y., and Lin, R. (2013). *Arabidopsis* Chromatin Remodeling Factor PICKLE Interacts with Transcription Factor HY5 to Regulate Hypocotyl Cell Elongation. The Plant Cell 25, 242–256. 10.1105/tpc.112.105742.

27. Zhang, D., Jing, Y., Jiang, Z., and Lin, R. (2014). The Chromatin-Remodeling Factor PICKLE Integrates Brassinosteroid and Gibberellin Signaling during Skotomorphogenic Growth in Arabidopsis. Plant Cell 26, 2472–2485. 10.1105/tpc.113.121848.

28. Xu, M., Hu, T., Smith, M.R., and Poethig, R.S. (2016). Epigenetic Regulation of Vegetative Phase Change in Arabidopsis. The Plant Cell 28, 28–41. 10.1105/tpc.15.00854.

29. Park, J., Oh, D.-H., Dassanayake, M., Nguyen, K.T., Ogas, J., Choi, G., and Sun, T. (2017). Gibberellin Signaling Requires Chromatin Remodeler PICKLE to Promote Vegetative Growth and Phase Transitions. Plant Physiol. 173, 1463–1474. 10.1104/pp.16.01471.

30. Liang, Z., Yuan, L., Xiong, X., Hao, Y., Song, X., Zhu, T., Yu, Y., Fu, W., Lei, Y., Xu, J., et al. (2022). The transcriptional repressors VAL1 and VAL2 mediate genome-wide recruitment of the CHD3 chromatin remodeler PICKLE in Arabidopsis. The Plant Cell 34, 3915–3935. 10.1093/plcell/koac217.

31. Song, M., Wang, Y., Ma, T., Terzaghi, W., He, K., Wang, H., Xu, B., Jing, Y., Lin, R., Deng, X.W., et al. (2025). PKL mediates H3K4me2 modification and spatial gene congregation in chromatin regulation. Nucleic Acids Res 53, gkaf1376. 10.1093/nar/gkaf1376.

32. Zha, P., Liu, S., Li, Y., Ma, T., Yang, L., Jing, Y., and Lin, R. (2020). The Evening Complex and the Chromatin-Remodeling Factor PICKLE Coordinately Control Seed Dormancy by Directly Repressing DOG1 in Arabidopsis. Plant Commun 1, 100011. 10.1016/j.xplc.2019.100011.

33. Liang, Z., Zhu, T., Yu, Y., Wu, C., Huang, Y., Hao, Y., Song, X., Fu, W., Yuan, L., Cui, Y., et al. (2024). PICKLE-mediated nucleosome condensing drives H3K27me3 spreading for the inheritance of Polycomb memory during differentiation. Molecular Cell 84, 3438–3454.e8. 10.1016/j.molcel.2024.08.018.

34. Fu, X., Li, C., Liang, Q., Zhou, Y., He, H., and Fan, L.-M. (2016). CHD3 chromatin-remodeling factor PICKLE regulates floral transition partially via modulating LEAFY expression at the chromatin level in Arabidopsis. Sci China Life Sci 59, 516–528. 10.1007/s11427-016-5021-x.

35. Yang, R., Zheng, Z., Chen, Q., Yang, L., Huang, H., Miki, D., Wu, W., Zeng, L., Liu, J., Zhou, J.-X., et al. (2017). The developmental regulator PKL is required to maintain correct DNA methylation patterns at RNA-directed DNA methylation loci. Genome Biol 18, 103. 10.1186/s13059-017-1226-y.

36. Zhang, X., Clarenz, O., Cokus, S., Bernatavichute, Y.V., Pellegrini, M., Goodrich, J., and Jacobsen, S.E. (2007). Whole-Genome Analysis of Histone H3 Lysine 27 Trimethylation in Arabidopsis. PLoS Biol 5, e129. 10.1371/journal.pbio.0050129.

37. Roudier, F., Ahmed, I., Bérard, C., Sarazin, A., Mary-Huard, T., Cortijo, S., Bouyer, D., Caillieux, E., Duvernois-Berthet, E., Al-Shikhley, L., et al. (2011). Integrative epigenomic mapping defines four main chromatin states in Arabidopsis: Organization of the Arabidopsis epigenome. The EMBO Journal 30, 1928–1938. 10.1038/emboj.2011.103.

38. Zhao, L., Zhou, Q., He, L., Deng, L., Lozano-Duran, R., Li, G., and Zhu, J.-K. (2022). DNA methylation underpins the epigenomic landscape regulating genome transcription in Arabidopsis. Genome Biol 23, 197. 10.1186/s13059-022-02768-x.

39. Ho, K.K., Zhang, H., Golden, B.L., and Ogas, J. (2013). PICKLE is a CHD subfamily II ATP-dependent chromatin remodeling factor. Biochim Biophys Acta 1829, 199–210. 10.1016/j.bbagrm.2012.10.011.

40. Carter, B., Bishop, B., Ho, K.K., Huang, R., Jia, W., Zhang, H., Pascuzzi, P.E., Deal, R.B., and Ogas, J. (2018). The Chromatin Remodelers PKL and PIE1 Act in an Epigenetic Pathway That Determines H3K27me3 Homeostasis in Arabidopsis. Plant Cell 30, 1337–1352. 10.1105/tpc.17.00867.

41. Hu, T., Manuela, D., Hinsch, V., and Xu, M. (2022). PICKLE associates with histone deacetylase 9 to mediate vegetative phase change in *Arabidopsis*. New Phytologist 235, 1070– 1081. 10.1111/nph.18174.

42. Li, W., Zhang, X., Zhang, Q., Li, Q., Li, Y., Lv, Y., Liu, Y., Cao, Y., Wang, H., Chen, X., et al. (2024). PICKLE and HISTONE DEACETYLASE6 coordinately regulate genes and transposable elements in *Arabidopsis*. Plant Physiology 196, 1080–1094. 10.1093/plphys/kiae369.

43. Alendar, A., and Berns, A. (2021). Sentinels of chromatin: chromodomain helicase DNA-binding proteins in development and disease. Genes Dev. 35, 1403–1430. 10.1101/gad.348897.121.

44. Farnung, L., Ochmann, M., and Cramer, P. (2020). Nucleosome-CHD4 chromatin remodeler structure maps human disease mutations. eLife 9, e56178. 10.7554/eLife.56178.

45. Jing, Y., Yang, Z., Yang, R., Zhang, Y., Qiao, W., Zhou, Y., and Sun, J. (2023). PKL is stabilized by MMS21 to negatively regulate Arabidopsis drought tolerance through directly repressing AFL1 transcription. New Phytol 239, 920–935. 10.1111/nph.18972.

46. Rider, S.D., Henderson, J.T., Jerome, R.E., Edenberg, H.J., Romero-Severson, J., and Ogas, J. (2003). Coordinate repression of regulators of embryonic identity by *PICKLE* during germination in *Arabidopsis*. The Plant Journal 35, 33–43. 10.1046/j.1365-313X.2003.01783.x.

47. Shim, J.S., Kubota, A., and Imaizumi, T. (2017). Circadian Clock and Photoperiodic Flowering in Arabidopsis: CONSTANS Is a Hub for Signal Integration. Plant Physiol. 173, 5–15. 10.1104/pp.16.01327.

48. Zhang, X., Chen, Y., Wang, Z., Chen, Z., Gu, H., and Qu, L. (2007). Constitutive expression of *CIR1* (*RVE2*) affects several circadian-regulated processes and seed germination in Arabidopsis. The Plant Journal 51, 512–525. 10.1111/j.1365-313X.2007.03156.x.

49. Matías-Hernández, L., Aguilar-Jaramillo, A.E., Marín-González, E., Suárez-López, P., and Pelaz, S. (2014). RAV genes: regulation of floral induction and beyond. Annals of Botany 114, 1459–1470. 10.1093/aob/mcu069.

50. Cheng, X.-F., and Wang, Z.-Y. (2005). Overexpression of COL9, a CONSTANS-LIKE gene, delays flowering by reducing expression of CO and FT in Arabidopsis thaliana. The Plant Journal 43, 758–768. 10.1111/j.1365-313X.2005.02491.x.

51. Gan, E.-S., Xu, Y., Wong, J.-Y., Geraldine Goh, J., Sun, B., Wee, W.-Y., Huang, J., and Ito, T. (2014). Jumonji demethylases moderate precocious flowering at elevated temperature via regulation of FLC in Arabidopsis. Nat Commun 5, 5098. 10.1038/ncomms6098.

52. Devlin, P.F., Robson, P.R.H., Patel, S.R., Goosey, L., Sharrock, R.A., and Whitelam, G.C. (1999). Phytochrome D Acts in the Shade-Avoidance Syndrome in Arabidopsis by Controlling Elongation Growth and Flowering Time1. Plant Physiol 119, 909–916. 10.1104/pp.119.3.909.

53. Mathieu, J., Yant, L.J., Mürdter, F., Küttner, F., and Schmid, M. (2009). Repression of flowering by the miR172 target SMZ. PLoS Biol 7, e1000148. 10.1371/journal.pbio.1000148.

54. Hu, H., Tian, S., Xie, G., Liu, R., Wang, N., Li, S., He, Y., and Du, J. (2021). TEM1 combinatorially binds to *FLOWERING LOCUS T* and recruits a Polycomb factor to repress the floral transition in *Arabidopsis*. Proc. Natl. Acad. Sci. U.S.A. 118, e2103895118. 10.1073/pnas.2103895118.

55. Hu, Y., Zhou, L., Yang, Y., Zhang, W., Chen, Z., Li, X., Qian, Q., Kong, F., Li, Y., Liu, X., et al. (2021). The gibberellin signaling negative regulator RGA-LIKE3 promotes seed storage protein accumulation. Plant Physiology 185, 1697–1707. 10.1093/plphys/kiaa114.

56. Li, C., Zheng, L., Wang, X., Hu, Z., Zheng, Y., Chen, Q., Hao, X., Xiao, X., Wang, X., Wang, G., et al. (2019). Comprehensive expression analysis of Arabidopsis GA2-oxidase genes and their functional insights. Plant Science 285, 1–13. 10.1016/j.plantsci.2019.04.023.

57. Rosenzweig, R., Nillegoda, N.B., Mayer, M.P., and Bukau, B. (2019). The Hsp70 chaperone network. Nat Rev Mol Cell Biol 20, 665–680. 10.1038/s41580-019-0133-3.

58. Ichino, L., Boone, B.A., Strauskulage, L., Harris, C.J., Kaur, G., Gladstone, M.A., Tan, M., Feng, S., Jami-Alahmadi, Y., Duttke, S.H., et al. (2021). MBD5 and MBD6 couple DNA methylation to gene silencing through the J-domain protein SILENZIO. Science 372, 1434– 1439. 10.1126/science.abg6130.

59. Zhong, Y., Moghaddas Sani, H., Paudel, B.P., Low, J.K.K., Silva, A.P.G., Mueller, S., Deshpande, C., Panjikar, S., Reid, X.J., Bedward, M.J., et al. (2022). The role of auxiliary domains in modulating CHD4 activity suggests mechanistic commonality between enzyme families. Nat Commun 13, 7524. 10.1038/s41467-022-35002-0.

60. Tian, Y., Jia, Q., Li, M., Sia, Y., Hu, P., Chen, K., Li, M., Li, X., Xu, Z., Ma, L., et al. (2025). Regulation of DNA translocation of chromatin remodeler enzyme Chd1 by exit DNA unwrapping. life. metab. 4, loaf013. 10.1093/lifemeta/loaf013.

61. Yan, L., and Chen, Z. (2020). A Unifying Mechanism of DNA Translocation Underlying Chromatin Remodeling. Trends in Biochemical Sciences 45, 217–227. 10.1016/j.tibs.2019.09.002.

62. Armache, K.-J., Garlick, J.D., Canzio, D., Narlikar, G.J., and Kingston, R.E. (2011). Structural Basis of Silencing: Sir3 BAH Domain in Complex with a Nucleosome at 3.0 Å Resolution. Science 334, 977–982. 10.1126/science.1210915.

63. Kuo, A.J., Song, J., Cheung, P., Ishibe-Murakami, S., Yamazoe, S., Chen, J.K., Patel, D.J., and Gozani, O. (2012). The BAH domain of ORC1 links H4K20me2 to DNA replication licensing and Meier-Gorlin syndrome. Nature 484, 115–119. 10.1038/nature10956.

64. Du, J., Zhong, X., Bernatavichute, Y.V., Stroud, H., Feng, S., Caro, E., Vashisht, A.A., Terragni, J., Chin, H.G., Tu, A., et al. (2012). Dual Binding of Chromomethylase Domains to H3K9me2-Containing Nucleosomes Directs DNA Methylation in Plants. Cell 151, 167–180. 10.1016/j.cell.2012.07.034.

65. Zhao, D., Zhang, X., Guan, H., Xiong, X., Shi, X., Deng, H., and Li, H. (2016). The BAH domain of BAHD1 is a histone H3K27me3 reader. Protein Cell 7, 222–226. 10.1007/s13238-016-0243-z.

66. Yang, Z., Qian, S., Scheid, R.N., Lu, L., Chen, X., Liu, R., Du, X., Lv, X., Boersma, M.D., Scalf, M., et al. (2018). EBS is a bivalent histone reader that regulates floral phase transition in Arabidopsis. Nat Genet 50, 1247–1253. 10.1038/s41588-018-0187-8.

67. Zhao, S., Lu, J., Pan, B., Fan, H., Byrum, S.D., Xu, C., Kim, A., Guo, Y., Kanchi, K.L., Gong, W., et al. (2023). TNRC18 engages H3K9me3 to mediate silencing of endogenous retrotransposons. Nature 623, 633–642. 10.1038/s41586-023-06688-z.

68. Machida, S., Takizawa, Y., Ishimaru, M., Sugita, Y., Sekine, S., Nakayama, J., Wolf, M., and Kurumizaka, H. (2018). Structural Basis of Heterochromatin Formation by Human HP1. Molecular Cell 69, 385–397.e8. 10.1016/j.molcel.2017.12.011.

69. Poepsel, S., Kasinath, V., and Nogales, E. (2018). Cryo-EM structures of PRC2 simultaneously engaged with two functionally distinct nucleosomes. Nat Struct Mol Biol 25, 154–162. 10.1038/s41594-018-0023-y.

70. Stoddard, C.I., Feng, S., Campbell, M.G., Liu, W., Wang, H., Zhong, X., Bernatavichute, Y., Cheng, Y., Jacobsen, S.E., and Narlikar, G.J. (2019). A Nucleosome Bridging Mechanism for Activation of a Maintenance DNA Methyltransferase. Molecular Cell 73, 73–83.e6. 10.1016/j.molcel.2018.10.006.

71. Mansfield, R.E., Musselman, C.A., Kwan, A.H., Oliver, S.S., Garske, A.L., Davrazou, F., Denu, J.M., Kutateladze, T.G., and Mackay, J.P. (2011). Plant Homeodomain (PHD) Fingers of CHD4 Are Histone H3-binding Modules with Preference for Unmodified H3K4 and Methylated H3K9. Journal of Biological Chemistry 286, 11779–11791. 10.1074/jbc.M110.208207.

72. Musselman, C.A., Ramírez, J., Sims, J.K., Mansfield, R.E., Oliver, S.S., Denu, J.M., Mackay, J.P., Wade, P.A., Hagman, J., and Kutateladze, T.G. (2012). Bivalent recognition of nucleosomes by the tandem PHD fingers of the CHD4 ATPase is required for CHD4-mediated repression. Proc Natl Acad Sci U S A 109, 787–792. 10.1073/pnas.1113655109.

73. Nodelman, I.M., Das, S., Faustino, A.M., Fried, S.D., Bowman, G.D., and Armache, J.-P. (2022). Nucleosome recognition and DNA distortion by the Chd1 remodeler in a nucleotide-free state. Nat Struct Mol Biol 29, 121–129. 10.1038/s41594-021-00719-x.

74. Yuan, J., Chen, K., Zhang, W., and Chen, Z. (2022). Structure of human chromatin-remodelling PBAF complex bound to a nucleosome. Nature 605, 166–171. 10.1038/s41586-022-04658-5.

75. Sia, Y., Pan, H., Chen, K., and Chen, Z. (2025). Structural insights into chromatin remodeling by ISWI during active ATP hydrolysis. Science 388, eadu5654. 10.1126/science.adu5654.

76. Lusser, A., Urwin, D.L., and Kadonaga, J.T. (2005). Distinct activities of CHD1 and ACF in ATP-dependent chromatin assembly. Nat Struct Mol Biol 12, 160–166. 10.1038/nsmb884.

77. Zhang, Y., LeRoy, G., Seelig, H.P., Lane, W.S., and Reinberg, D. (1998). The dermatomyositis-specific autoantigen Mi2 is a component of a complex containing histone deacetylase and nucleosome remodeling activities. Cell 95, 279–289. 10.1016/s0092-8674(00)81758-4.

78. Low, J.K.K., Webb, S.R., Silva, A.P.G., Saathoff, H., Ryan, D.P., Torrado, M., Brofelth, M., Parker, B.L., Shepherd, N.E., and Mackay, J.P. (2016). CHD4 Is a Peripheral Component of the Nucleosome Remodeling and Deacetylase Complex. Journal of Biological Chemistry 291, 15853–15866. 10.1074/jbc.M115.707018.

79. Qiu, X.-B., Shao, Y.-M., Miao, S., and Wang, L. (2006). The diversity of the DnaJ/Hsp40 family, the crucial partners for Hsp70 chaperones. Cell. Mol. Life Sci. 63, 2560–2570. 10.1007/s00018-006-6192-6.

80. Kampinga, H.H., and Craig, E.A. (2010). The HSP70 chaperone machinery: J proteins as drivers of functional specificity. Nat Rev Mol Cell Biol 11, 579–592. 10.1038/nrm2941.

81. V. Rajan, V.B., and D’Silva, P. (2009). Arabidopsis thaliana J-class heat shock proteins: cellular stress sensors. Funct Integr Genomics 9, 433–446. 10.1007/s10142-009-0132-0.

82. Harris, C.J., Scheibe, M., Wongpalee, S.P., Liu, W., Cornett, E.M., Vaughan, R.M., Li, X., Chen, W., Xue, Y., Zhong, Z., et al. (2018). A DNA methylation reader complex that enhances gene transcription. Science 362, 1182–1186. 10.1126/science.aar7854.

83. Zhao, Q., Lin, R., Li, L., Chen, S., and He, X. (2019). A methylated-DNA-binding complex required for plant development mediates transcriptional activation of promoter methylated genes. JIPB 61, 120–139. 10.1111/jipb.12767.

84. Zheng, S.-Y., Guan, B.-B., Yuan, D.-Y., Zhao, Q.-Q., Ge, W., Tan, L.-M., Chen, S.-S., Li, L., Chen, S., Xu, R.-M., et al. (2023). Dual roles of the Arabidopsis PEAT complex in histone H2A deubiquitination and H4K5 acetylation. Molecular Plant 16, 1847–1865. 10.1016/j.molp.2023.10.006.

85. Luo, Y., Hou, X., Zhang, C., Tan, L., Shao, C., Lin, R., Su, Y., Cai, X., Li, L., Chen, S., et al. (2020). A plant-specific SWR1 chromatin-remodeling complex couples histone H2A.Z deposition with nucleosome sliding. The EMBO Journal 39, e102008. 10.15252/embj.2019102008.

86. Tan, L.-M., Liu, R., Gu, B.-W., Zhang, C.-J., Luo, J., Guo, J., Wang, Y., Chen, L., Du, X., Li, S., et al. (2020). Dual Recognition of H3K4me3 and DNA by the ISWI Component ARID5 Regulates the Floral Transition in Arabidopsis. Plant Cell 32, 2178–2195. 10.1105/tpc.19.00944.

87. Kim, D., Paggi, J.M., Park, C., Bennett, C., and Salzberg, S.L. (2019). Graph-based genome alignment and genotyping with HISAT2 and HISAT-genotype. Nat Biotechnol 37, 907–915. 10.1038/s41587-019-0201-4.

88. Robinson, M.D., McCarthy, D.J., and Smyth, G.K. (2010). edgeR: a Bioconductor package for differential expression analysis of digital gene expression data. Bioinformatics 26, 139–140. 10.1093/bioinformatics/btp616.

89. Wu, C.-J., Yuan, D.-Y., Liu, Z.-Z., Xu, X., Wei, L., Cai, X.-W., Su, Y.-N., Li, L., Chen, S., and He, X.-J. (2023). Conserved and plant-specific histone acetyltransferase complexes cooperate to regulate gene transcription and plant development. Nat. Plants 9, 442–459. 10.1038/s41477-023-01359-3.

90. Zhang, Y., Liu, T., Meyer, C.A., Eeckhoute, J., Johnson, D.S., Bernstein, B.E., Nusbaum, C., Myers, R.M., Brown, M., Li, W., et al. (2008). Model-based Analysis of ChIP-Seq (MACS). Genome Biol 9, R137. 10.1186/gb-2008-9-9-r137.

91. Zang, C., Schones, D.E., Zeng, C., Cui, K., Zhao, K., and Peng, W. (2009). A clustering approach for identification of enriched domains from histone modification ChIP-Seq data. Bioinformatics 25, 1952–1958. 10.1093/bioinformatics/btp340.

92. Guo, J., Cai, G., Li, Y.-Q., Zhang, Y.-X., Su, Y.-N., Yuan, D.-Y., Zhang, Z.-C., Liu, Z.-Z., Cai, X.-W., Guo, J., et al. (2022). Comprehensive characterization of three classes of Arabidopsis SWI/SNF chromatin remodelling complexes. Nat. Plants 8, 1423–1439. 10.1038/s41477-022-01282-z.

93. Langmead, B., and Salzberg, S.L. (2012). Fast gapped-read alignment with Bowtie 2. Nat Methods 9, 357–359. 10.1038/nmeth.1923.

94. Stark, R., and Brown, G. DiffBind : differential binding analysis of ChIP-Seq peak data.

95. Qian, F., Zhao, Q., Zhang, T., Li, Y., Su, Y., Li, L., Sui, J., Chen, S., and He, X. (2021). A histone H3K27me3 reader cooperates with a family of PHD finger-containing proteins to regulate flowering time in *Arabidopsis*. JIPB 63, 787–802. 10.1111/jipb.13067.

96. Dyer, P.N., Edayathumangalam, R.S., White, C.L., Bao, Y., Chakravarthy, S., Muthurajan, U.M., and Luger, K. (2003). Reconstitution of Nucleosome Core Particles from Recombinant Histones and DNA. In Methods in Enzymology (Elsevier), pp. 23–44. 10.1016/S0076-6879(03)75002-2.

97. Yan, L., Wang, L., Tian, Y., Xia, X., and Chen, Z. (2016). Structure and regulation of the chromatin remodeller ISWI. Nature 540, 466–469. 10.1038/nature20590.

98. Ye, Y., Wu, H., Chen, K., Clapier, C.R., Verma, N., Zhang, W., Deng, H., Cairns, B.R., Gao, N., and Chen, Z. (2019). Structure of the RSC complex bound to the nucleosome. Science 366, 838–843. 10.1126/science.aay0033.

99. Liu, X., Li, M., Xia, X., Li, X., and Chen, Z. (2017). Mechanism of chromatin remodelling revealed by the Snf2-nucleosome structure. Nature 544, 440–445. 10.1038/nature22036.

