## Supplemental Figures and Tables for "J-domain proteins stimulate PKL-mediated chromatin compaction at H3K4-hypomethylated genomic loci"

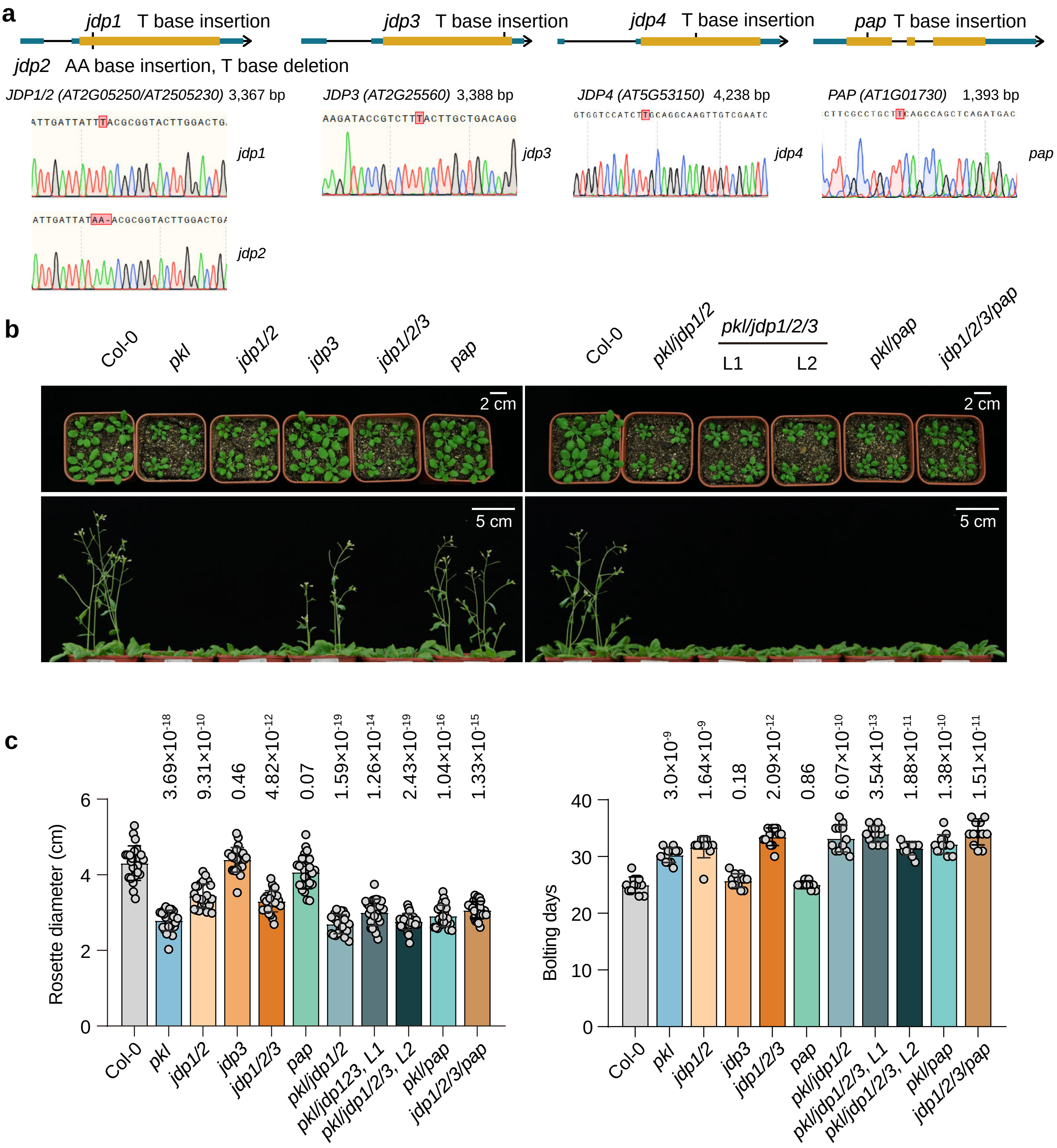

**Extended Data Fig. 1 | Mutation sites of Arabidopsis mutant plants used in the study.**
(**a**) Schematic of mutated genes and sequencing validation of target mutation regions. For each panel, the top illustrates the schematic of mutated genes, with mutation sites marked; the bottom displays sequencing snapshots, where the mutated nucleotide positions highlighted in red. In the schematic of mutated genes, orange blocks denote coding regions, and blue blocks denote untranslated regions. Mutation sites of the *jdp1/2* , *jdp3*, *jdp4* and *pap* mutant lines are shown individually. (**b,c**) Morphological phenotypes (**b**) and quantitative analysis (**c**) of plants at 3 weeks (upper panel) and 5 weeks (lower panel) post-germination. Rosette width was measured at the pre-bolting stage. Data are presented as mean ± SD, with individual data points denoted by circles. *p* value was determined using a two-tailed Student's t-test.

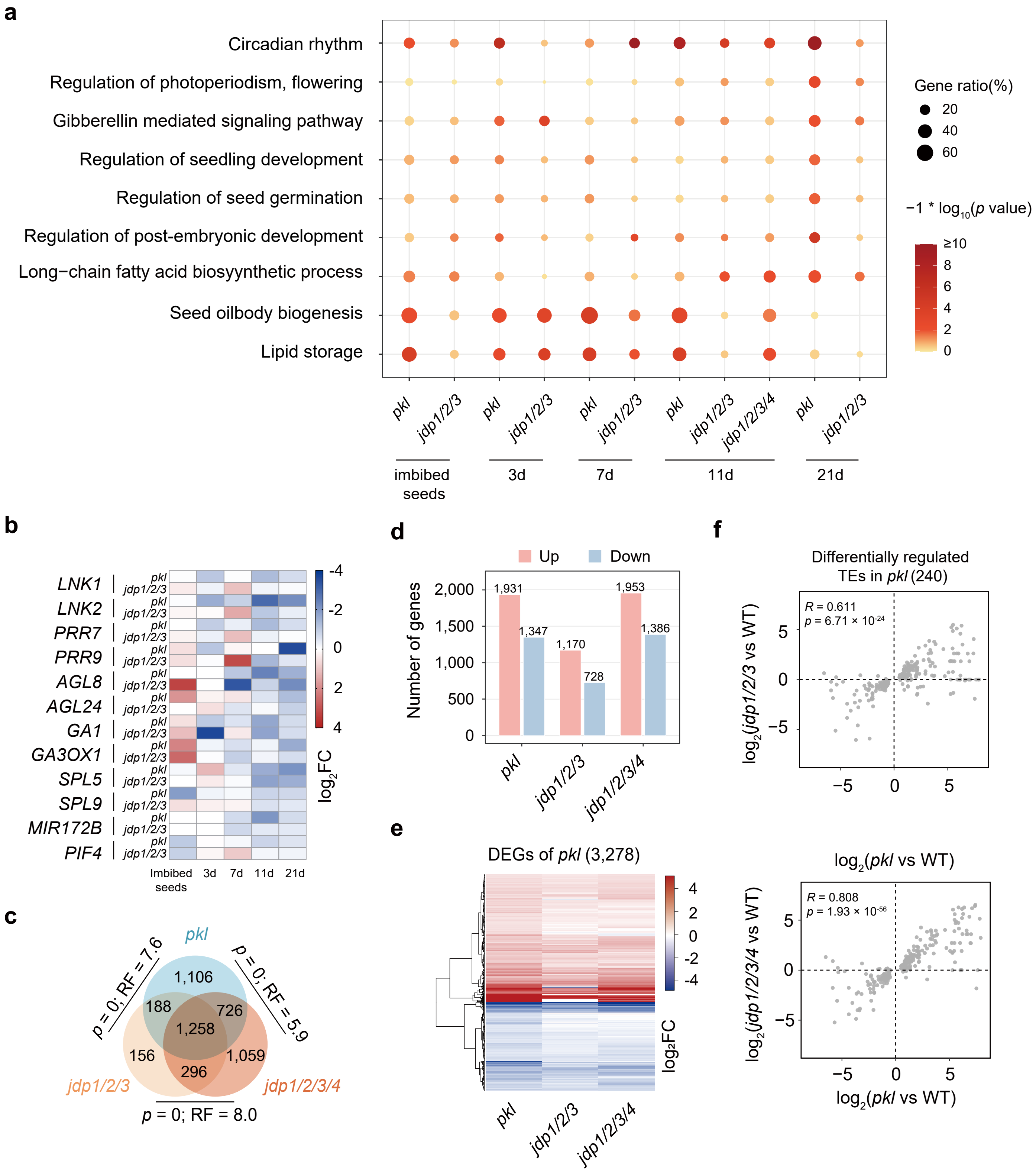

**Extended Data Fig. 2 | Analysis of differentially expressed genes and TEs in *pkl*, *jdp1/2/3*, and *jdp1/2/3/4* mutants.** (**a**) GO enrichment analyses were conducted separately for the DEGs related to flowering and embryonic development identified in *pkl* and *jdp* mutants relative to the Col-0 wild-type control in different phases. The y-axis represents enriched GO biological process terms. Bubble size indicates the proportion of DEGs annotated to each term and the color gradient represents the *p* value. (**b**) Heatmap showing the $\log_2$FC of positive regulators in flowering in *pkl* and *jdp1/2/3* mutants. The $\log_2$FC values are represented by color bars. (**c**) Overlap of DEGs identified in *pkl*, *jdp1/2/3*, and *jdp1/2/3/4* mutants. The significance of overlaps is indicated by *p* values and representation factors (RF). *p* values were determined using a one-tailed hypergeometric test. (**d**) Number of upregulated and downregulated DEGs in *pkl*, *jdp1/2/3*, and *jdp1/2/3/4* mutants relative to Col-0 wild-type plants. (**e**) Heatmaps displaying the expression changes of *pkl*, *jdp1/2/3*, and *jdp1/2/3/4* mutants relative to wild-type plants. Analyses were performed using DEGs identified in the *pkl* mutant (|$\log_2$ fold change ($\log_2$FC)| > 0.5; false discovery rate (FDR) < 0.05). The $\log_2$FC values are represented by color bars. (**f**) Correlation of TE expression changes caused by *pkl*, and *jdp1/2/3/4* mutations. Analyses were performed using differentially expressed TEs identified in the *pkl* mutant (|$\log_2$FC| > 0.5; FDR < 0.05).

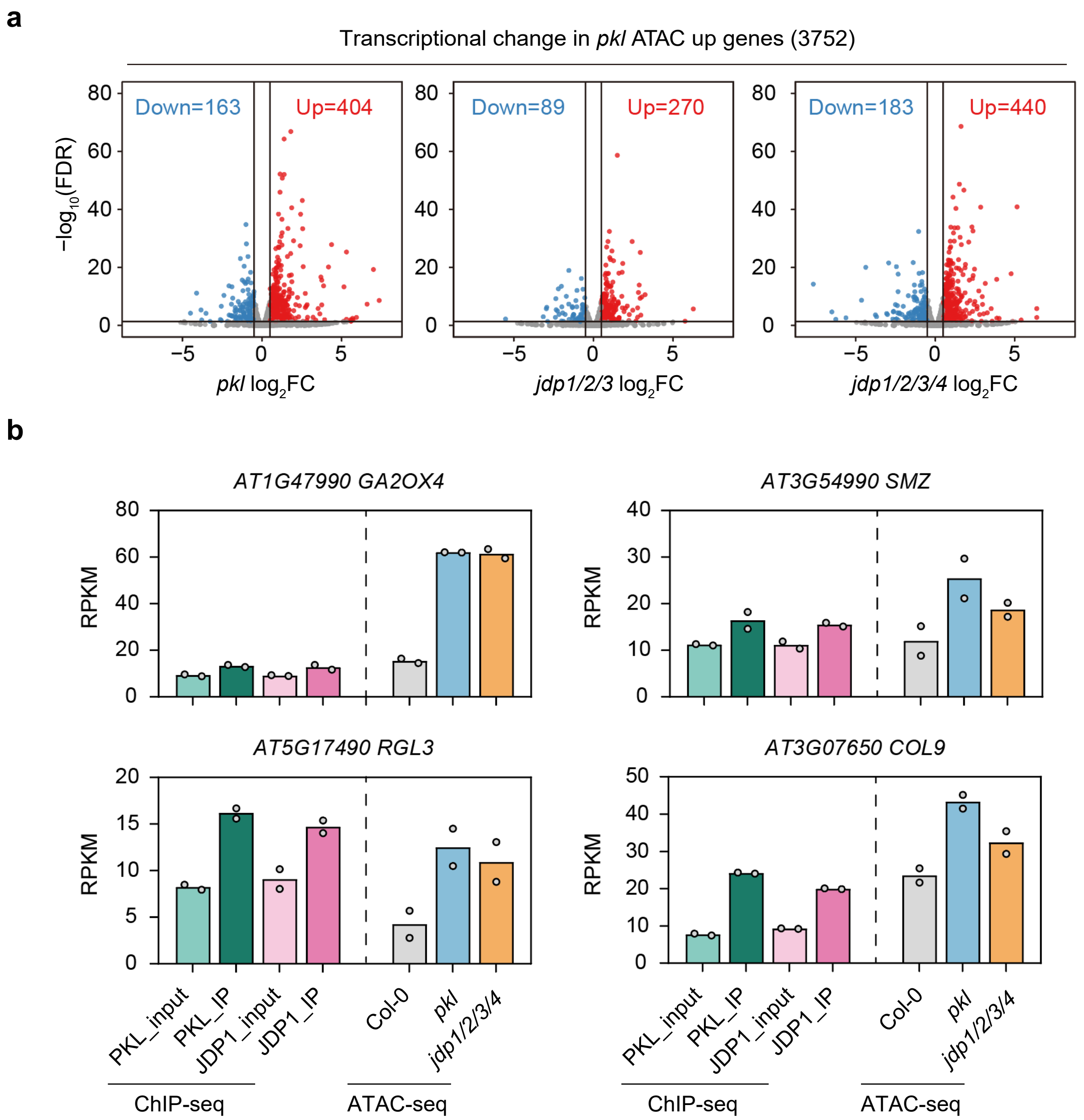

**Extended Data Fig. 3 | Effects of *pkl*, *jdp1/2/3*, and *jdp1/2/3/4* mutations on gene expression and chro-matin accessibility.**

(**a**) Volcano plots exhibiting transcriptional changes of genes with increased accessiblity of *pkl* in *pkl*, *jdp1/2/3* and *jdp1/2/3/4* mutants compared to wild type. X-axis shows $\log_2$ fold change between mutants and wild type ($\log_2$FC), and Y-axis shows -$\log_{10}$ false discovery rate (FDR). (**b**) Quantitative analysis of PKL and JDP1 enrichment peaks and increased chromatin accessibility peaks in *pkl* and *jdp1/2/3/4* mutants within the 2 kb region upstream of the TSS for representative flowering repressor genes.

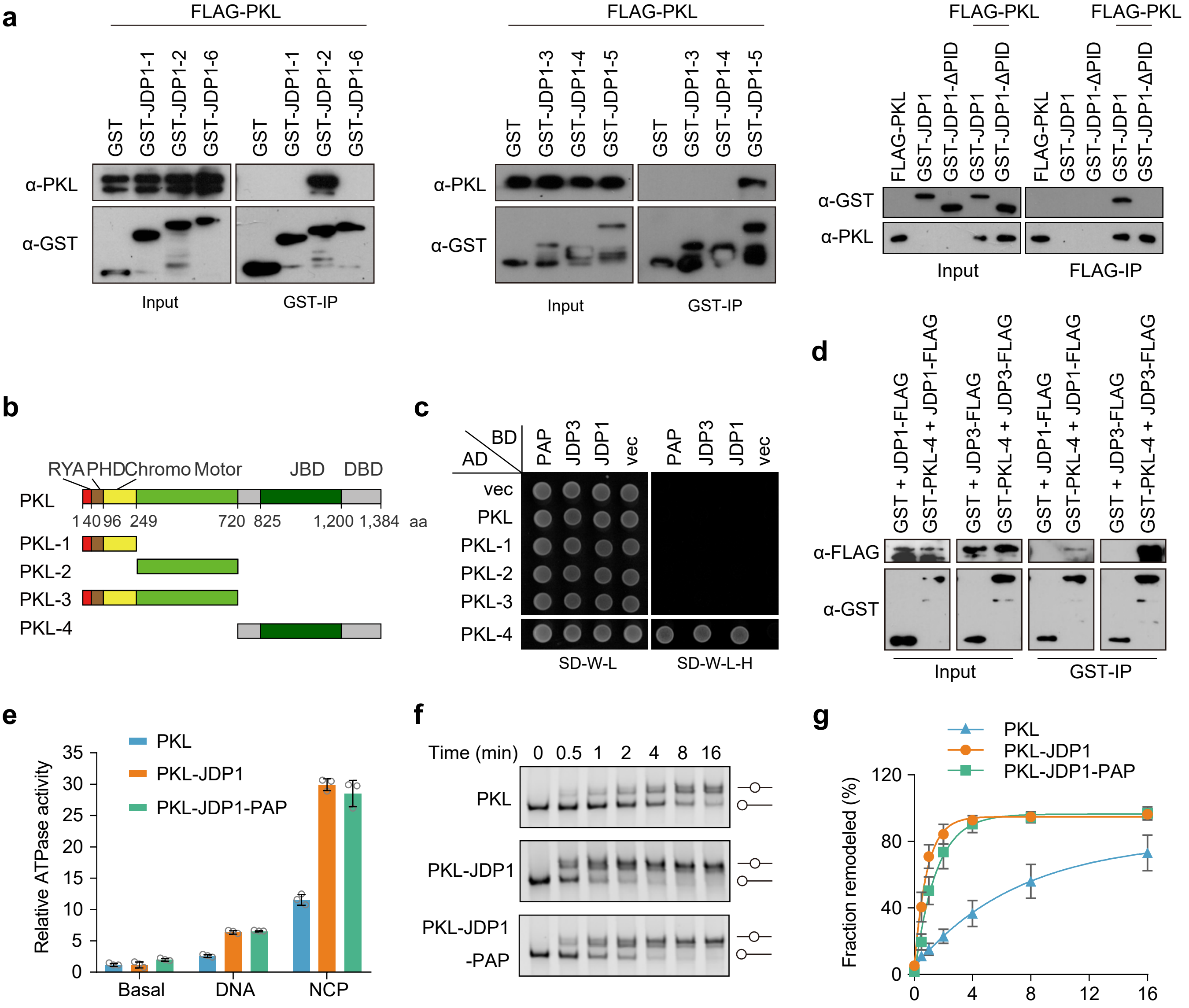

**Extended Data Fig. 4 | Characterization of the PKL-JDP interaction and effects of JDP1 and PAP on PKL ATPase and remodeling activities.**

(**a**) *In vitro* pull-down assays characterizing the interaction between PKL and truncated JDP1 protein variants. The truncated JDP1 constructs utilized in these experiments are illustrated in Fig. 3A. (**b**) Schematic representation of full-length and truncated PKL variants used for yeast two-hybrid and pull-down assays. Domains of PKL characterized in the present study and prior reports are indicated. (**c**) Yeast two-hybrid assays to determine the interaction of JDP1, JDP3, and PAP with truncated PKL variants. (**d**) Validation of the interaction between the PKL truncated variant PKL-4 and JDP1 or JDP3 by pull-down assays. (**e**) Relative ATPase activities of PKL, PKL-JDP1 and PKL-JDP1-PAP. Data are presented as mean ± SD (n = 3). (**f** and **g**) Chromatin-remodeling activities of PKL, PKL-JDP1 and PKL-JDP1-PAP toward the 0N40 nucleosome. Representative gels are presented in (**f**), with quantification of the fractions of nucleosome slide shown in (**g**). Data are presented as mean ± SD (n = 3). The initial reaction rates ($min^{-1}$) estimated by fitting and data are 0.11, 1.07 and 0.68 for PKL, PKL-JDP1, and PKL-JDP1-PAP, respectively.

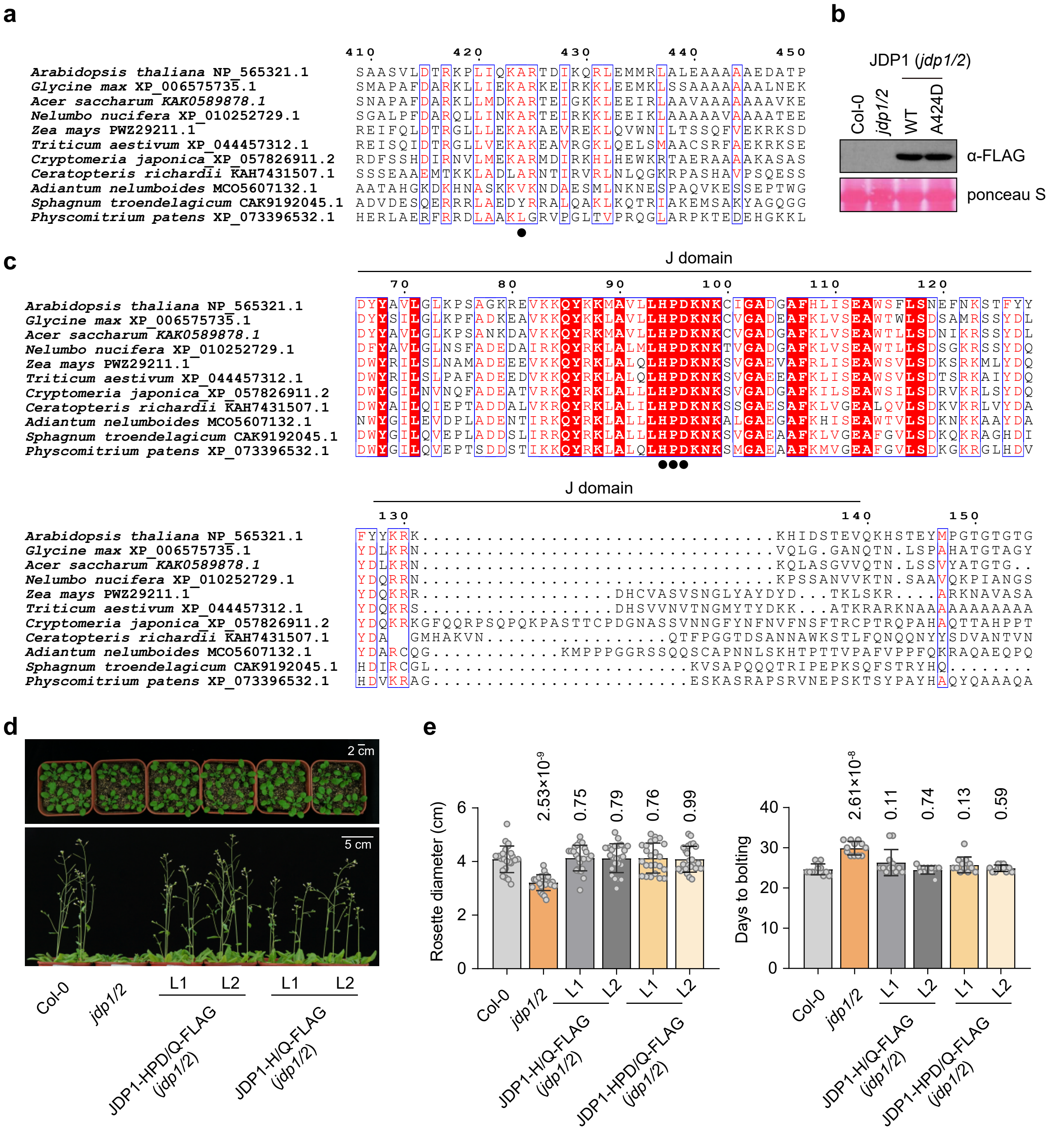

**Extended Data Fig. 5 | Sequence and functional characterization of the PID domain and J domain in JDP1.** (**a**) Sequence alignment of the α-helix of PID domain from Arabidopsis JDP1 and its homologs in other plant lineages. Protein sequences were retrieved via BLAST searches of the NCBI database and aligned using CLUSTALW. Residue A424 of JDP1 is marked with a dot. (**b**) Immunoblot analysis of FLAG-tagged JDP1-WT and JDP1-A424D variant protein abundance in the *jdp1/2* mutant background. Ponceau S staining is shown as a loading control. (**c**) Sequence alignment of the J domain from Arabidopsis JDP1 and its homologs in other plant species. Protein sequences were acquired via BLAST searches of the NCBI database and aligned using CLUSTALW. The conserved HPD motif, which is essential for HSP70 interaction, is marked with dots. (**d**) Morphological phenotypes of plants at three weeks (top) and five weeks (bottom) post-germination. (**e**) Quantification of rosette width at the pre-bolting stage (left) and days to bolting (right). Individual biological replicates are represented by circles. Data are presented as mean ± SD. *p* values were determined by a two-tailed Student's t-test.

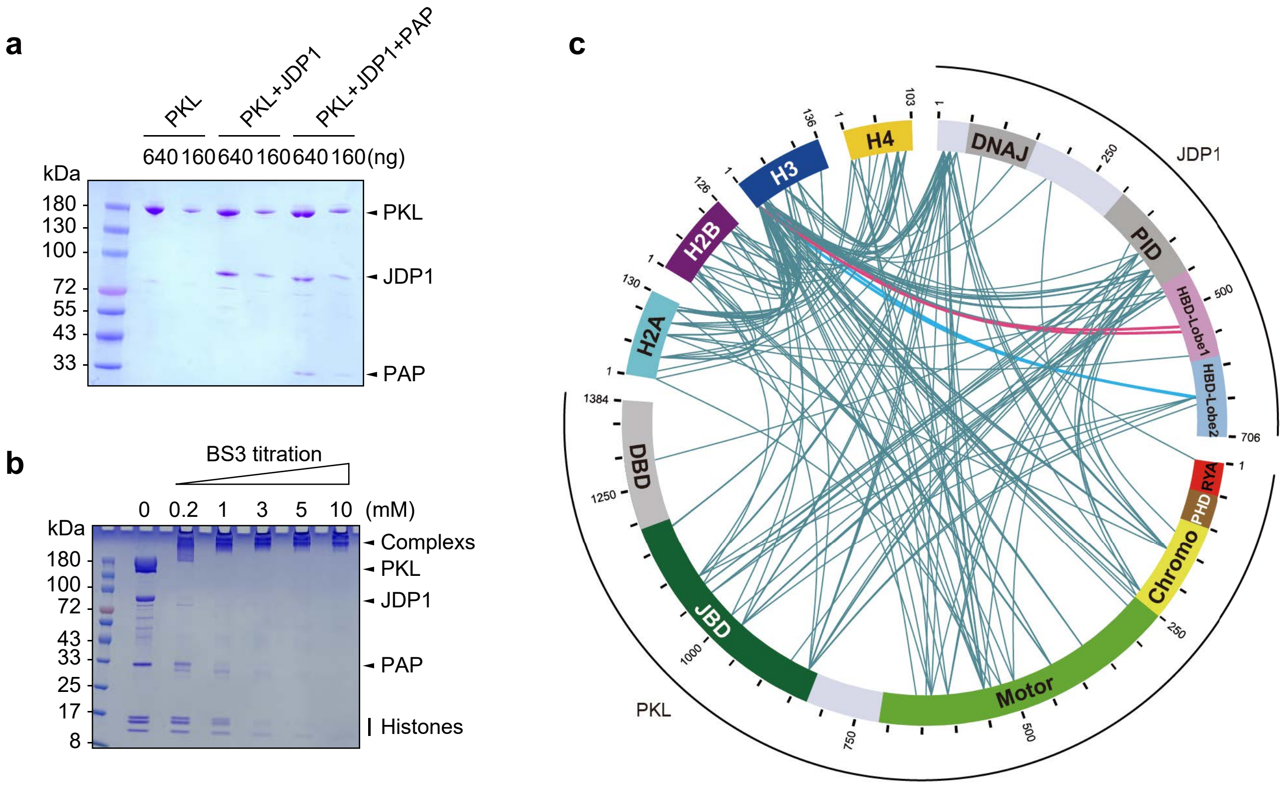

**Extended Data Fig. 6 | Cross-linking mass spectrometry analysis of the PKL-JDP1 complex bound to the nucleosomes.**

(**a**) Coomassie Brilliant Blue staining of Strep-tagged PKL, His-tagged JDP1 and His-tagged PAP proteins. (**b**, **c**) Cross-linking mass spectrometry analysis (CL-MS) to determine inter-molecular interactions of the PKL-JDP-Nucleosome Core Particle (NCP) complex. Coomassie Brilliant Blue staining of cross-linked protein mixtures (**b**) and schematic representation of identified intermolecular cross-linking sites (**c**) are shown. Domains are colored as in Fig. 3A and fig. S5B. The results support the JBD-PID and CTD-H3 interactions.

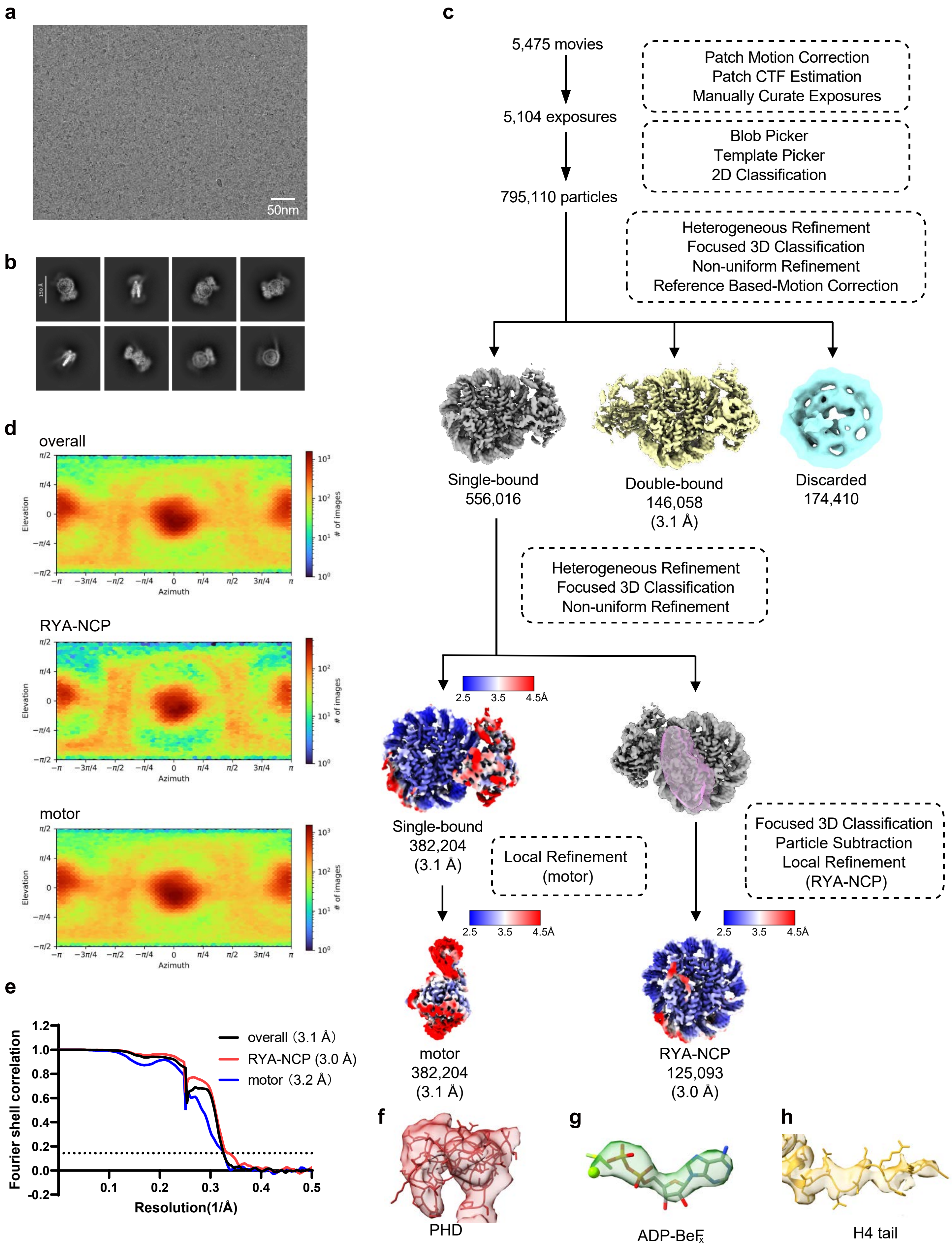

**Extended Data Fig. 7 | Workflow of cryo-EM data processing.**

(**a**) Representative cryo-EM micrograph. (**b**) Representative 2D classes of selected particles. (**c**) Flowchart of the cryo-EM data processing. The overall resolution of the complex was improved to 3.1 Å. Some weak density appeared at a low contour level on the H2A-H2B surface of the nucleosome, indicating RYA motif of PKL interacts with the acidic patch of H2A-H2B. After focused 3D classification and local refinement with a soft mask, RYA-NCP part was refined to 3.0 Å. (**d**) Angular distribution of cryo-EM particles in the final round of refinement. (**e**) Gold-standard Fourier shell correlation (FSC) curves, showing the overall nominal resolutions of 3.1 Å, 3.0 Å and 3.1 Å for the overall complex, RYA-NCP and motor, respectively. (**f**) Local density map of the PHD domain of PKL. (**g**) Local density map of the ADP-BeFx bound by PKL. (**h**) Local density map of the H4 tail bound by PKL.

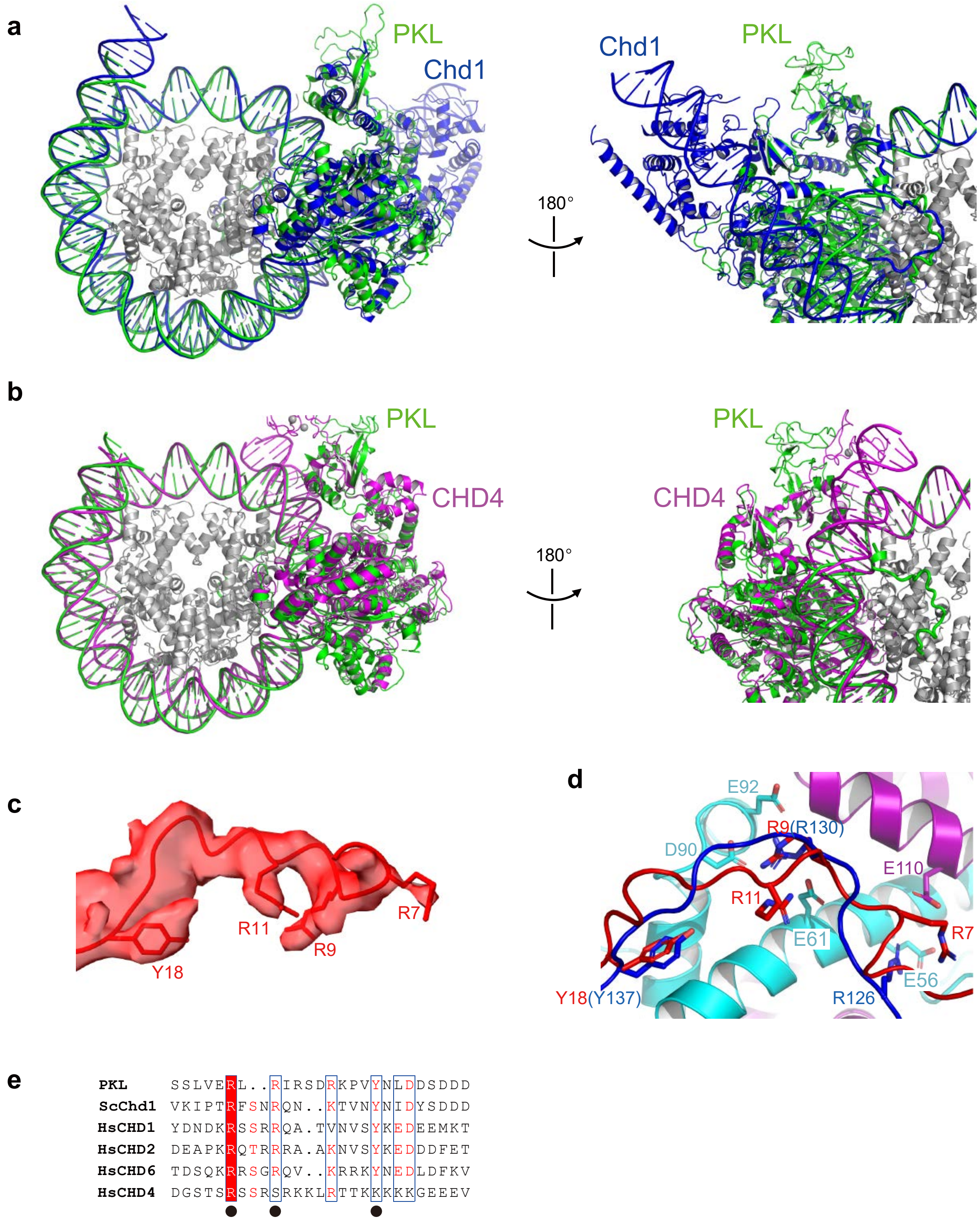

**Extended Data Fig. 8 | Structural comparison of PKL with other CHD family proteins,.**
(**a**) Structural comparison of PKL (colored green) and yeast Chd1 (colored blue, PDB code 7TN2) bound to the nucleosome. The histone octamers are aligned. (**b**) Structural comparison of PKL and human CHD4 (colored magenta, PDB code 6RYR) bound to the nucleosome. The histone octamers are aligned. (**c**) Local cryo-EM map around the RYA motif of PKL. (**d**) Structural comparison of the RYA motifs from PKL (colored red) and Chd1 (colored blue, PDB code 7TN2) in H2A-H2B binding. (**e**) Multiple sequence alignments of CHD family proteins. Conserved residues of the RYA motif are highlighted with dots. The RYA motif is absent from CHD4.

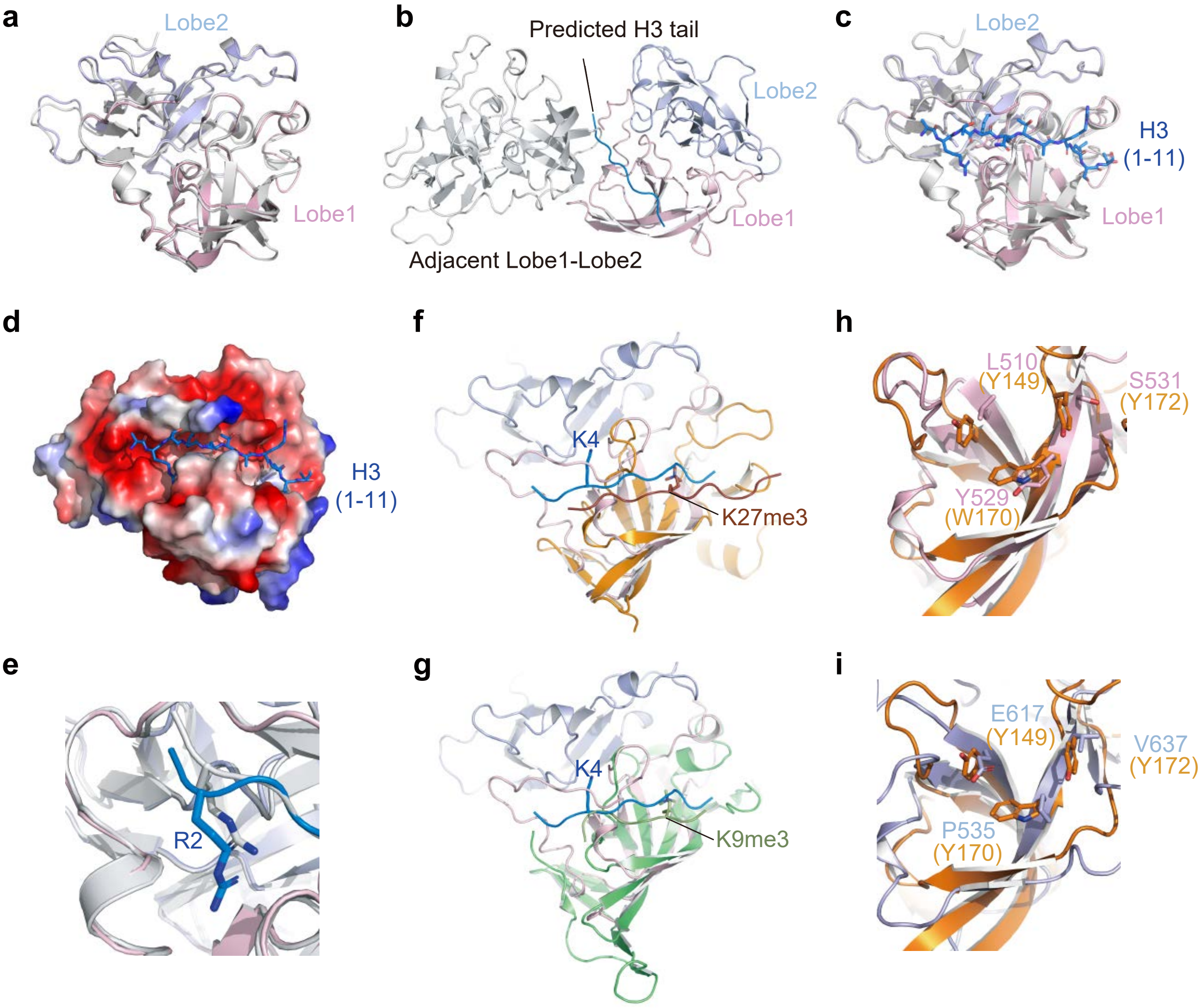

**Extended Data Fig. 9 | Analysis of the H3-binding ability of the HBD in JDP1.**
(**a**) Structural comparison of crystal structure of Lobe1-Lobe2 (colored pink and light blue, respectively) and predicted Lobe1-Lobe2 (colored grey). (**b**) The predicted H3-binding pocket is blocked by the crystal contacts in the crystals of HBD. (**c**) Structural comparison of crystal structure of Lobe1-Lobe2 (colored pink and light blue, respectively) of JDP1 bound to H3 peptide (colored blue) and Lobe1-Lobe2 alone (colored grey). (**d**) Crystal structure of JDP1-Lobe1-Lobe2 bound to H3 peptide. JDP1-Lobe1-Lobe2 is shown in surface representation, with electrostatic potential calculated using PyMOL. Red, negative electrostatic potential; blue, positive electrostatic potential. (**e**) Structural comparison of H3R2 observed in the crystal structure (colored blue) and predicted by AlphaFold3 (colored grey). (**f**) Structural comparison of the HBD of JDP1 bound to the H3 peptide (colored blue) and the BAH domain of AIPP3 bound to H3K27me3 peptide (colored orange, PDB code 7CCE). (**g**) Structural comparison of the HBD of JDP1 bound to H3 and the BAH domain of TNRC18 bound to H3K9me3 peptide (colored green, PDB code 8DS8). (**h**, **i**) Structural comparison of the Lobe1 (**h**) and Lobe2 (**i**) domains of JDP1 with the BAH domain of AIPP3, respectively.

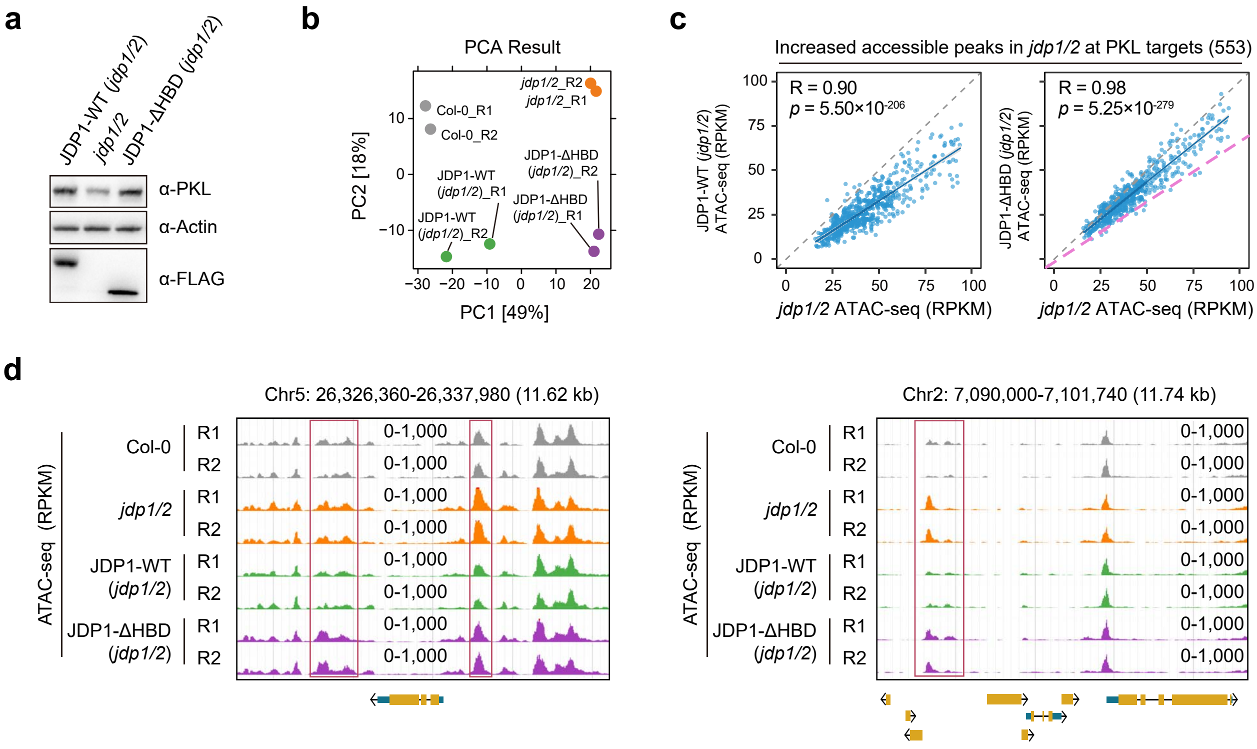

**Extended Data Fig. 10 | Effects of HBD truncation on JDP1 chromatin association and on JDP1-dependent restriction of chromatin accessibility.**

(**a**) Immunoblot analysis of FLAG-tagged JDP1-WT and JDP1-ΔHBD variant protein abundance in the *jdp1/2* mutant background. (**b**) Principal component analysis (PCA) of ATAC-seq data from Col-0 wild type, *jdp1/2* mutants, and complementation lines expressing JDP1-WT and JDP1-ΔHBD. Percentages indicate the variance explained by PC1 and PC2. (**c**) Scatter plots showing reduction of ATAC-seq enrichment mediated by JDP1-WT and JDP1-ΔHBD in the *jdp1/2* background. Analysis was performed on 553 PKL target loci with increased chromatin accessibility in *jdp1/2* relative to Col-0 wild type (FDR < 0.05). Fitting lines were shown in blue. In the right panel, the pink dashed line indicates the fitting line from the left panel for reference. (**d**) Genome browser view of ATAC-seq signals in Col-0, *jdp1/2*, and complementation lines expressing JDP1-WT and JDP1-ΔHBD.

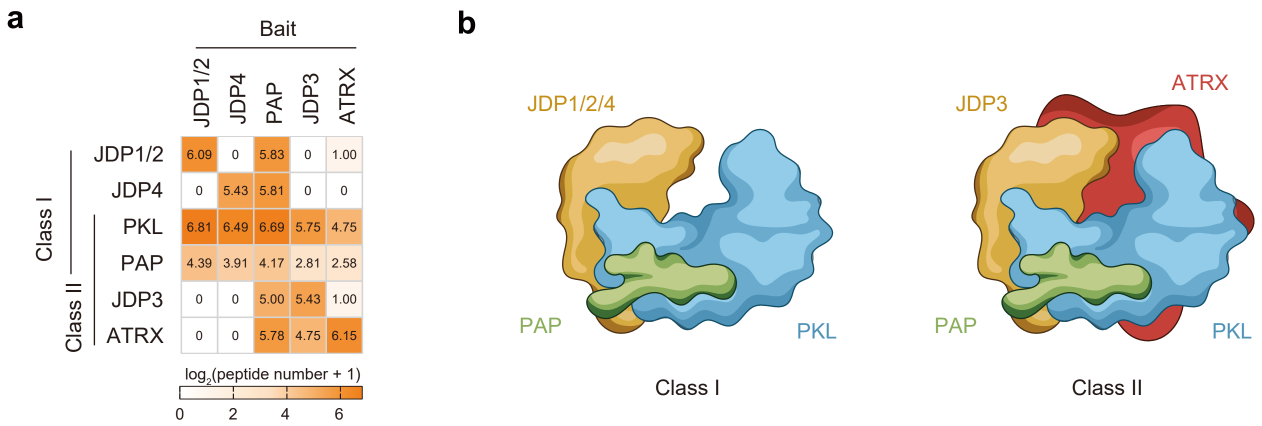

**Supplementary Fig. 1 | PKL–JDP interactions and structural organization of PKL–JDP complexes.**
(**a**) Heatmap showing interactions among PKL, JDPs, PAP and ATRX proteins. The abundance of co-precipitated proteins was quantified based on peptide counts obtained from AP-MS data. (**b**) Schematic representation of two classes of PKL–JDP complexes, based on AlphaFold3 predictions.

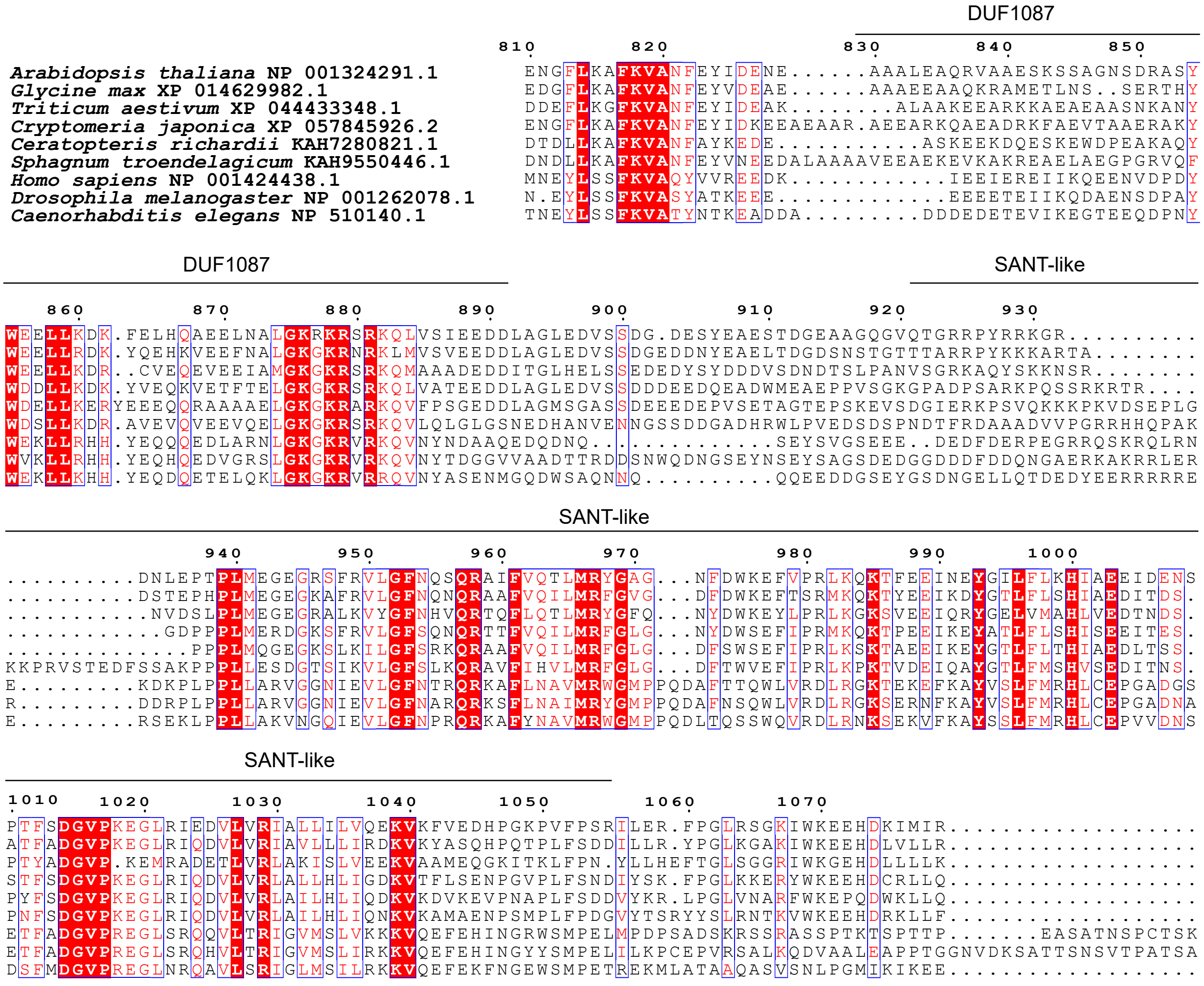

**Supplementary Fig. 2 | Sequence alignment of JDP-binding domain of PKL and its homologs.** Protein sequences were retrieved via BLAST searches of the NCBI database and aligned using CLUSTALW (https://www.genome.jp/tools-bin/clustalw). Conserved DUF1087 and SANT-like domains are indicated.

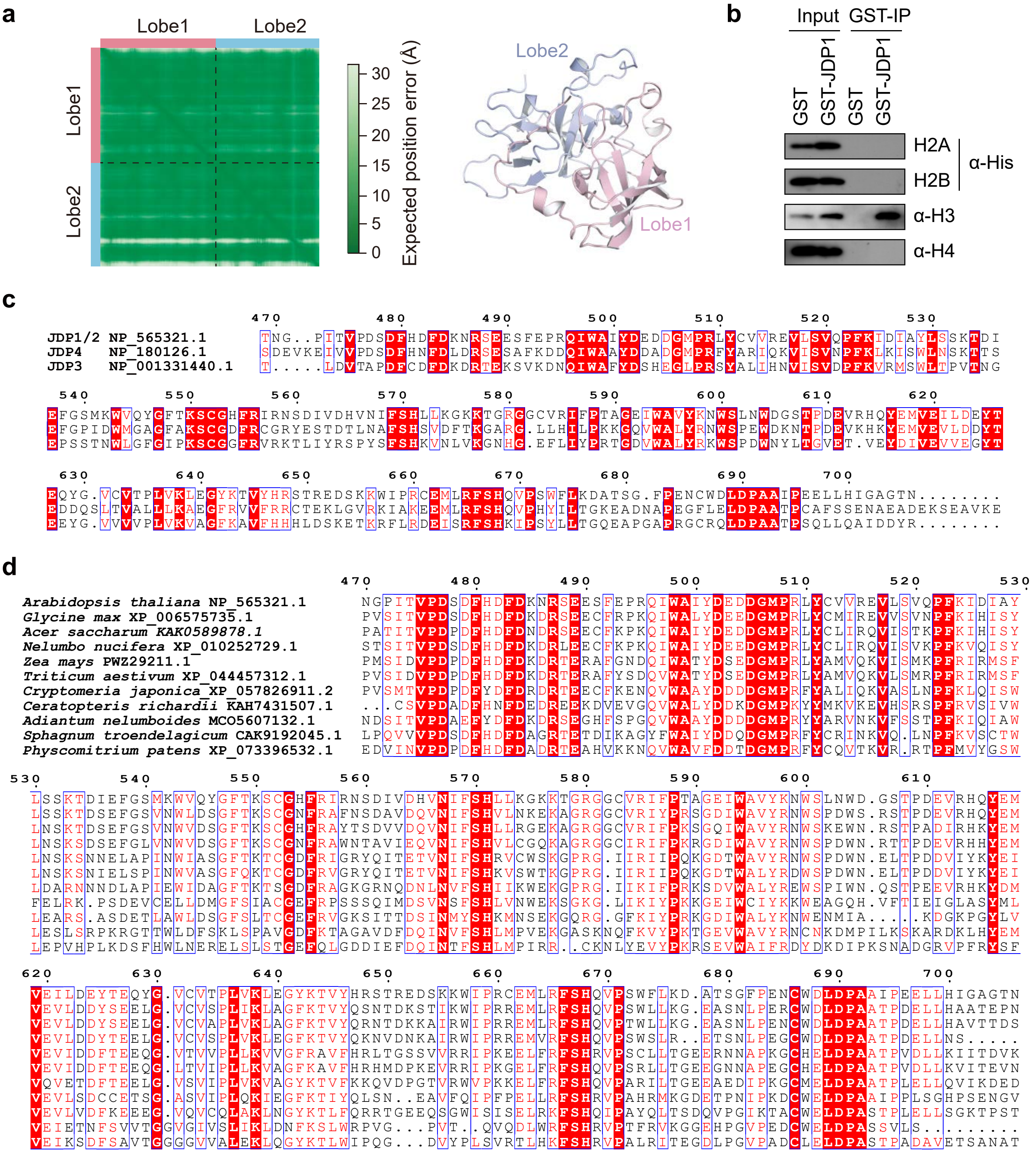

**Supplementary Fig. 3 | Sequence analysis and functional characterization of the HBD.**
(**a**) PAE matrix (left) and the corresponding structure (right) of the predicted HBD, with Lobe1 and Lobe2 indicated. (**b**) Determination of the interaction between JDP1 and H2A, H2B, H3, or H4 by pull-down assays. GST-tagged JDP1 was individually incubated with each His-tagged histone for pull-down assays, followed by immunoblot analysis. (**c**) Sequence alignment of the HBD domain of JDP1/2, JDP3 and JDP4. Protein sequences were retrieved from the NCBI database, and aligned using CLUSTALW. (**d**) Sequence alignment of the HBD domain in plant JDP1 homologs. Sequences were obtained via BLAST searches in the NCBI database, and aligned using CLUSTALW.

**Supplementary Table 1**. Cryo-EM data collection, refinement and validation statistics

| | PKL-NCP (EMD-68253, PDB 22FZ) | RYA-NCP (EMD-68244) | PKL-motor (EMD-83227) |
|---|---|---|---|
| **Data collection and processing** | | | |
| Microscope | Titan Krios | Titan Krios | Titan Krios |
| Camera | K3 | K3 | K3 |
| Magnification (nominal) | 29,000 | 29,000 | 29,000 |
| Electron exposure (e-/Å2) | 50 | 50 | 50 |
| Number of frames collected | 32 | 32 | 32 |
| Automation software | AutoEMation2 | AutoEMation2 | AutoEMation2 |
| Voltage (kV) | 300 | 300 | 300 |
| Micrographs (no.) | 5,475 | 5,475 | 5,475 |
| Defocus range (μm) | -1.2— -1.8 | -1.2— -1.8 | -1.2— -1.8 |
| Pixel size (Å) | 0.485 | 0.485 | 0.485 |
| Symmetry imposed | C1 | C1 | C1 |
| Initial particle images (no.) | 795,110 | 795,110 | 795,110 |
| Final particle images (no.) | 382,204 | 125,093 | 382,204 |
| Map resolution (Å)(masked) | 3.1 | 3.0 | 3.1 |
| FSC threshold | 0.143 | 0.143 | 0.143 |
| Map sharpening *B* factor (Å2) | | -40 | |
| **Refinement** | | | |
| Initial model used(pdb mode) | | | |
| Refinement package | Phenix | | |
| Model-map scores | | | |
| CC(mask) | 0.84 | | |
| CC(peaks) | 0.79 | | |
| CC(volume) | 0.84 | | |
| R.m.s deviations | | | |
| Bond lengths (Å) | 0.006 | | |
| Bond angle (°) | 0.611 | | |
| C-beta deviation | 0.00 | | |
| EMRinger score | 2.42 | | |
| CaBLAM outliers | 1.74 | | |
| **Validation** | | | |
| MolProbity score | 1.79 | | |
| Clashscore | 9.09 | | |
| Poor rotamers (%) | 0.00 | | |
| Ramachandran plot | | | |
| Favored (%) | 95.55 | | |
| Allowed (%) | 4.39 | | |
| Disallowed (%) | 0.06 | | |

**Supplementary Table 2.** Data collection and refinement statistics (molecular replacement)

| | HBD (PDB 22FI) | HBD-H3 (PDB 22EZ) |
|---|---|---|
| **Data collection** | | |
| Space group | $P3_121$ | C121 |
| Cell dimensions | | |
| a, b, c (Å) | 85.82, 85.82, 79.60 | 63.31, 73,84, 63,45 |
| α, β, γ (°) | 90, 90, 120 | 90.00, 115.10, 90.00 |
| Resolution (Å)* | 54-2.12 (2.23-2.12) | 50-2.24 (2.30-2.24) |
| $R_{sym}$ / $R_{merge}$ | 0.216 (2.951) | 0.054 (0.176) |
| $I/\sigma I$ | 11.4 (1.9) | 34.8 (10.5) |
| Completeness (%) | 96.5 (100.0) | 99.5 (99.8) |
| Redundancy | 16.1 (13.9) | 6.4 (6.3) |
| CC1/2 | 0.99 (0.49) | 1.00 (0.97) |
| **Refinement** | | |
| Resolution (Å) | 37-2.12 | 45-2.24 |
| No. reflections | 18824 | 12300 |
| $R_{work}$ / $R_{free}$ | 23.1/26.8 | 20.1/23.4 |
| No. atoms | | |
| Protein | 1910 | 1979 |
| Ligand/ion | 1 | / |
| Water | 130 | 92 |
| *B*-factors | | |
| Protein | 45.2 | 42.9 |
| Ligand/ion | 31.2 | / |
| Water | 44.7 | 40.7 |
| R.m.s. deviations | | |
| Bond lengths (Å) | 0.003 | 0.009 |
| Bond angle (°) | 0.58 | 0.95 |

*Values in parentheses are for highest-resolution shell.
